# The transcriptomic signature of age and sex is not conserved in human primary myocytes

**DOI:** 10.1101/2024.12.17.629041

**Authors:** Séverine Lamon, Megan Soria, Ross Williams, Annabel Critchlow, Karel Van Belleghem, Andrew Garnham, Akriti Varshney, Traude Beillharz, Danielle Hiam, Mark Ziemann

**Affiliations:** Institute for Physical Activity and Nutrition, School of Exercise and Nutrition Sciences, Deakin University, Geelong, Australia; School of Life and Environmental Sciences, Deakin University, Geelong, Australia; Development and Stem Cells Program, Monash Biomedicine Discovery Institute and Department of Biochemistry and Molecular Biology, Monash University, Melbourne, Australia; The Novo Nordisk Foundation Centre for Stem Cell Medicine (reNEW), Murdoch Children’s Research Institute, Melbourne 3052, Victoria, Australia; Bioinformatics Working Group, Burnet Institute, Melbourne, Australia

**Keywords:** Aging, Sex, Transcriptome, Muscle Cells, Primary Cell Culture

## Abstract

**Background:** Human primary muscle cell (HPMC) lines derived from skeletal muscle biopsies are potentially powerful tools to interrogate the molecular pathways underlying fundamental muscle mechanisms. HPMCs retain their genome in culture, but many endogenous circulating factors are not present in the *in vitro* environment, or at concentrations that do not mirror physiological levels. To address the assumption that HPMCs are valid models of age and sex-specificity in human muscle research, we examined to what extent HPMC lines retain their source phenotype in culture.

**Methods:** Biopsies from the v*astus lateralis* muscle were collected from ten males aged 18-30, ten females aged 18-30 and ten males aged 60-75 recruited from a general, healthy population. A portion of the muscle was used for the establishment of 30 individual HMPC lines. The remaining sample was immediately snap frozen and stored for further analysis. RNA was extracted from muscle tissue samples and their corresponding, fully differentiated HMPCs and analysed using RNA Sequencing. To compare their transcriptomic signature, principal component analysis (PCA), differential expression analysis, single-cell deconvolution and pathway enrichment analysis were conducted in *R*.

**Results:** A comparison of the transcriptomic signature of 30 human muscle biopsies and their corresponding HPMCs indicated a near-complete lack of retention of the genes and pathways differentially regulated *in vivo* when compared to their *in vitro* equivalent, with the exception of several genes encoded on the Y-chromosome.

**Conclusions:** The diversity of resident cell populations in muscle tissue and the lack of sex- and age-dependent circulating factors in the cellular milieu likely contribute to these observations, which call for caution when using HPMCs as an experimental model of human muscle sex or age.

## Introduction

Skeletal muscle is essential to human posture, movement and metabolism and is a critical component of healthy ageing (1). Primary muscle cell lines (2) established from human or rodent muscle tissue are potentially powerful tools to interrogate the molecular pathways underlying fundamental muscle mechanisms. The phenotype of a cell line can be modified using molecular tools that may or may not alter its genotype, which reduces the number of animals used in research and is cheaper and faster than animal experiments. Primary muscle cell lines provide an alternative to the traditional C_2_C_12_ line, an immortalized murine myoblast-derived cell line (3) with cycles of ‘active’ to ‘resting’ that closely resembles those of satellite cells that cycle between an activated and a quiescent state (4). While complex to establish (2), more expensive to maintain and limited in the number of times they can be passaged, human primary muscle cell (HPMC) lines may be molecularly and metabolically closer to human physiology (5) despite their intrinsic genetic heterogeneity.

Skeletal muscle is a sex-biased tissue. We and others have shown that it has between 60 and 3000 genes that are differentially expressed in males and females, depending on the technique utilized and statistical power of each study (6–10). This contributes to numerous sex-specific phenotypic traits, for example in muscle growth, regenerative capacity (11) and fibre type distribution (12) (see (13) for a full review). It follows that many fundamental mechanisms underlying skeletal muscle biology are sex-specific and cannot be inferred from single- or combined-sex study designs. Skeletal muscle is also profoundly affected by ageing, which facilitates an irregular yet progressive series of molecular and cellular dysfunctions that inevitably impact muscle structure, function and metabolism (1, 14–19). A common, directly observable consequence is age-related muscle loss; an insidious, progressive process characterized by 3–8% reduction in muscle mass and function per decade from the fifth decade (20). At the transcriptome level, recent studies highlight the pathways deregulated during ageing (21, 22), which were largely similar in males and females despite sex differences in individual gene expression (23) and magnitude of the changes (24).

HPMC lines retain the genome of their donor in culture, including some of the differences stemming from the sex chromosome complement (25, 26), with primary differences in gene expression and DNA methylation (27). Primary cultures allow tight control of the micro-environment of cells. Many endogenous circulating factors, including sex- and age-dependent factors such as steroid hormones and growth factors, are however either not present in the *in vitro* environment or at concentrations that do not mirror physiological levels. In addition, whole muscle biopsies not only contain myoblasts and mature muscle cells, but also fibro-adipogenic progenitors, immune cells, fibroblasts and endothelial cells (28–30). These sub-populations are filtered out when preparing human primary cultures, introducing another source of variability. Whether and to what extent HPMC lines maintain their source phenotypes in culture is therefore in need of full examination. For example, we and others showed that some of the ‘ageing’ phenotypes of the myocytes derived from old subjects are lost in cell culture models (31–33), while other age- and metabolism-related traits that are partially conserved may include lipolysis, insulin sensitivity (34) and DNA methylation (35). The same applies to male and female derived cell lines (27). While some sex differences may be conserved in HPMCs, the extent of the sex differences mediated by gonadal hormones that is retained in culture is unknown.

Regardless of these limitations, HPMCs are commonly used to probe age and sex-specific muscle molecular pathways in the muscle literature. To address the assumption that HPMCs are valid models of age and sex-specificity, we conducted the first large-scale assessment of the transcriptome of young and older male and female cell lines and compared them to the muscle transcriptome of their donor. We identified a distinct transcriptomic signature of age and sex in human muscle tissue, which, except for a few Y-encoded genes, was not conserved in their corresponding HMPCs. Understanding to what extent sex- and age-specific differences mediated by internal and external factors are retained in culture is essential to establish cause-and-effect relationships between molecular regulators and age- and sex-specific phenotypes before conducting validation studies.

## Methods

### Ethics

This research study was granted ethical approval by the Deakin University Human Research Ethics Committee (DUHREC 2021-307). All participants provided written, informed consent before taking part in the study, which was conducted in accordance with the Declaration of Helsinki (36) and its later amendments.

### Participants

Participants were recruited from the local community as part of the broader FAMe (37) and MALe studies, which include males and females participants aged 18-80 (ethics approval number: DUHREC 2021-307). Consent, recruitment and experimental protocols are described in details elsewhere (37). Ten males aged 18-30, ten females aged 18-30 and ten males aged 60-75 were selected to form the sub-cohort described in this study using a randomization function. Inclusion criteria included biologically male and female participants with a normal body mass index (BMI) of 18-35kg/m^2^ and deemed generally healthy through completion of a medical questionnaire and assessment by a medical practitioner. Exclusion criteria included type 2 diabetes, a current diagnosis or < 5 years of remission from cancer, a current diagnosis of neurological or neuromuscular disorder, the presence of any infectious bloodborne disease, allergies to anaesthetics, current status as a smoker, the current use of any dietary supplements with the potential to have an effect on muscle metabolism (e.g. thyroxine), any intellectual or mental impairment making it difficult to understand the invasiveness of the medical procedures and provide proper informed consent, and any history of gender affirming treatment of surgery, or hormonal treatment excluding hormonal contraceptives (females only). Measures of moderate-to-vigorous physical activity, dietary protein intake, muscle strength (via leg press 1RM) and appendicular lean mass index (ALMI) were conducted as described in (37) prior to sample collection. Characteristics of the sub-cohort of participants used in this study are presented in Table 1.

**Table 1.** Characteristics of the sub-cohort of participants selected for this study.

| **Group** |  | **Males (young)**<br>**N = 10** | **Males (older)**<br>**N = 10** | **Females**<br>**(young) N = 10** |
| --- | --- | --- | --- | --- |
| **Age**<br>**(year)** | **Mean** | 23.22 | 67.50 | 26.90 |
|  | **SD** | 3.15 | 4.19 | 6.51 |
| **Weight**<br>**(kg)** | **Mean** | 76.25 | 80.56 | 72.33 |
|  | **SD** | 10.94 | 13.46 | 13.69 |
| **Height**<br>**(cm)** | **Mean** | 177.03 | 175.04 | 165.73 |
|  | **SD** | 7.22 | 7.15 | 7.92 |
| **BMI**<br>**(kg.m<sup>2</sup>)** | **Mean** | 24.26 | 26.21 | 26.31 |
|  | **SD** | 2.66 | 3.26 | 4.38 |
| **MVPA**<br>**(min.day<sup>-1</sup>)** | **Mean** | 45.67 | 19.79 | 41.20 |
|  | **SD** | 28.79 | 16.03 | 28.00 |
| **Protein intake**<br>**(g.kg<sup>-1</sup>.day<sup>-1</sup>)** | **Mean** | 1.96 | 1.40 | 1.70 |
|  | **SD** | 0.83 | 0.52 | 0.69 |
| **Leg press 1RM**<br>**(kg)** | **Mean** | 312.01 | 233.85 | 107.80 |
|  | **SD** | 66.45 | 44.05 | 29.22 |
| **ALMI**<br>**(kg.m<sup>-2</sup>)** | **Mean** | 8.90 | 8.41 | 7.50 |
|  | **SD** | 0.74 | 0.96 | 1.21 |

BMI: body mass index; MVPA: moderate-to-vigorous physical activity; RM, repetition maximum; ALMI: appendicular lean mass index.

### Hormonal status

Female participants were pre-menopausal, not currently pregnant or lactating and completed a menstrual cycle history questionnaire prior to entering the study. Muscle sample collection aimed to occur between day 1-7 of the menstrual cycle, when the ratio of oestrogen to progesterone is the lowest to minimise fluctuations in sex hormones as a potential confounding variable. A blood sample was collected on the same day to control and measures of oestrogen, progesterone, LH and FSH measured by high-performance liquid chromatography mass spectrometry (Triple Quad 5500, Sciex, Framingham, MA) at Monash Health Pathology laboratory (Victoria, Australia) were used to confirm the menstrual status according to (38).

### Muscle sample collection

Participants reported to the laboratory in the morning in a fasted state from 9pm the previous evening. They were provided with a standardized meal of pasta and tomato sauce to eat the night before, and had abstained from caffeine, alcohol, and vigorous physical activity in the 24 hours prior to the visit (37).

A muscle biopsy was collected from the *vastus lateralis* muscle using the percutaneous needle technique (39) modified to include suction (40) as described in our previous work (17). First, the skin was be sterilised and anaesthetised with 1% Lidocaine without epinephrine. Following this, an incision was made through the skin and fascia and a muscle sample of ∼ 100-200 mg was excised. Fifty – 100 mg of muscle was placed in ice-cold serum free Hams F10 nutrient mixture (Thermo Fisher Scientific) for the establishment of HPMC lines. The remaining sample was immediately snap frozen in liquid isopentane and stored in liquid nitrogen for further analysis.

### Human primary cell lines

Thirty individual primary muscle cell lines were established in as previously described in detail by our group in (33). An extra purification step was added when the cells had reached 70-80% confluence, and myogenic satellite cells were purified using Magnetic Activated Cell Sorting (MACS) with anti-CD56+ microbeads Miltenyi Biotec) as previously described (41, 42).

Myoblasts were maintained in proliferation media made up of Ham’s F-10 medium (Thermo Fisher Scientific) containing 20% FBS, 25 ng/mL fibroblast growth factor (bFGF; Promega), 1% penicillin-streptomycin, and 0.5% amphoteromycin (Life Technologies). At approximately 70–80% confluence, proliferation medium was replaced with differentiation medium made up of DMEM (Life Technologies) containing 5.5mM glucose, 2% HS (New Zealand origin; Gibco, Life Technologies) and 1% penicillin/streptomycin. Differentiation medium was replaced every 48 h. RNA collection was performed on the 9^th^ day of differentiation. The presence of mycoplasma was tested using *contam* (https://github.com/markziemann/contam/tree/main) and revealed an average of 17.8 reads per million (RPM) (range: 8.0 – 22.5 RPM) per sample aligning to one of five antibiotic-resistant mycoplasmic genomes. As a result, the cells were deemed mycoplasma free (43).

### RNA extraction

RNA was extracted from 10 mg samples of snap frozen *vastus lateralis* tissue or from the pooled content of three technical replicates from 6-well plates, using the AllPrep DNA/RNA/miRNA Universal kit (Qiagen) according to the manufacturer’s instructions including proteinase K and DNase treatment. RNA quality and quantity were assessed using the TapeStation System according to manufacturer’s instructions (Agilent Technologies). An RNA integrity number (RIN) of >7 was considered acceptable for downstream RNA sequencing analysis.

### RNA sequencing, transcript quantification and quality control

The RNAseq libraries were prepared using the Illumina TruSeq Stranded Total RNA with Ribo-Zero Gold protocol and sequenced with 150-bp paired-end reads on the Illumina Novaseq6000 (Macrogen Oceania Platform). Paired-end reads were trimmed of adapters with an average phred score below 10 using Skewer (44). Reads were aligned using Kallisto (45) and used to map reads to the reference human transcriptome from Ensembl (genome assembly GRCh38, release-111) and to generate transcript counts. The quality of the reads before and after trimming and alignment was monitored using FastQC (v0.11.9) (46) and MultiQC (v1.14) (47). The resulting transcriptome count matrix was then mapped to its corresponding gene using biomaRt package (version 2.56.1) (48) and were filtered out of low counts (each gene should contain at least an average of 10 counts across all samples) resulting into 17,789 Ensembl genes IDs for use as input for downstream analysis.

### Principal Component Analysis

Filtered count matrices were normalised via the Variance Stabilising Transformation function from the DESeq2 package (v1.40.2) (49). PCA plots were generated for multiple comparisons using the RNAseqQC package (v0.1.4) (50).

### Differential expression and enrichment analysis

Differential expression (DE) analysis was performed with the DESeq2 package and pipeline using R (v4.3.2) and RStudio (Build 386, “Cherry Blossom” for Ubuntu). A threshold of 0.05 for the Benjamini and Hochberg (BH-adjusted) p values is set as a threshold for significance and a log_2_ fold change of 1 and -1 is the threshold set for upregulated and downregulated differentially expressed genes (DEGs) respectively.

Functional enrichment was performed using the GSEA function from the clusterProfiler package (v4.12.6) using ranked gene lists based on log□ fold changes from DESeq2 to capture intrinsic differences in cell populations between muscle tissue and cell lines. Reactome gene sets from *Homo sapiens* using the msigdbr package (v7.5.1) were then curated for skeletal muscle tissue activation using <u>PubChem</u> and <u>The Human Protein Atlas</u>, with all non-filtered gene sets presented in Supplementary Tables. The minimum gene set size was set to 10 and an FDR value cutoff of 0.05. Multidimensional enrichment analysis was conducted using *mitch*, which ranks genes by their differential expression based on DESeq2 test statistic before conducting MANOVA based enrichment test (51). Rank-rank gene density contour heatmaps were used to depict the similarity between gene expression profiles. Together, *mitch* and GSEA are designed to detect subtle expression changes across genes that may not be apparent at the individual gene level but, when considered collectively, can indicate pathway-level enrichment (51) despite the lack of widespread individual gene significance. *Mitch* outputs were curated for skeletal muscle tissue activation using <u>PubChem</u> and <u>The Human Protein Atlas</u>, with all non-filtered gene sets presented in Supplementary Tables.

### Differentiation time course

A differentiation time course was established based on RNASeq counts from our previous work (52). The data was deposited on the Gene Expression Ominbus (GEO) platform under accession number GSE168897(52). Briefly, one male and one female human skeletal muscle cell line were established according to our existing protocols (33). Myocytes were seeded in 6-well plates and allowed to differentiate as described above. Cell lines were harvested and RNA collected 24hrs after seeding (T1), at the onset of differentiation (T2), as well as after one (T3), three (T4), five (T5) and seven (T6) days of differentiation. Two technical replicates per time point per cell line were analysed, with representative immunohistochemistry images available elsewhere (52).

For initial data preparation, a minimum count threshold of 10 was applied across samples to filter low-expression genes or genes not reliably detected across samples, resulting in 15,859 Ensembl gene IDs. This global filtering approach improves the sensitivity of detecting true DEGs by reducing noise and statistical burden. It ensures that only robustly expressed genes are retained, strengthening the reliability of downstream analyses such as gene set enrichment and pathway analysis (53). Normalization was conducted through variance-stabilizing transformation (VST), implemented using the *vst* function from DESeq2 (49) ensuring comparable expression data suitable for downstream analysis. Pairwise differential expression analysis was performed across timepoints, and further analysis using the Likelihood Ratio Test in the DESeq2 package was used to identify how genes are differentially expressed between individual timepoints. Genes with similar patterns of expression were clustered using the degPatterns function from the DEGreport package (v1.40.1) (http://lpantano.github.io/DEGreport/).

### Differentiation score calculations

Principal component analysis (PCA) was performed on the normalized and transposed count matrix using the *prcomp* function from the *stats* package in R (v4.4.1). The first principal component (PC1) loadings were extracted to capture the most significant variance across genes related to differentiation. Using these PC1 loadings, a weighted sum was computed for each sample, wherein each gene’s normalized expression was weighted by its loading in PC1, providing a summary differentiation score per sample based on gene expression. The resulting weighted sums were then min-max scaled, converting values between 0 and 1, enabling consistent comparison across samples and time points. For each sample, the scaled differentiation scores were aggregated by time point, averaging the scores for each time point to represent the differentiation progression over time.

### Application of Differentiation Scores to Cell RNASeq Data

To apply differentiation scores to cell-level data, the normalized cell counts were subset to retain only genes present in the differentiation dataset. The cell differentiation scores were classified into discrete time points by applying threshold values derived from the time-point scores in the differentiation dataset. This classification assigned each cell differentiation score to a specific time point (e.g., T1 through T6), facilitating a temporal interpretation of differentiation states. Scores surpassing the defined thresholds were assigned a classification of “T6_plus” for cells exhibiting differentiation levels beyond the maximum time point threshold.

### Deconvolution of bulk muscle tissues to infer cell type proportion

To explore the proportion of different cell types within muscle tissue samples, deconvolution of the bulk skeletal muscle transcriptome data was performed using the MuSiC (Multi-subject Single Cell deconvolution) package in R (54). Publicly available single-cell RNA-seq data from the Tabula Sapiens atlas, specifically the skeletal muscle dataset (“TS_Muscle.h5ad” which is available via FigShare), was used as a reference. Preprocessing of the single-cell data was conducted using Scanpy in Python (v3.12). Cells with fewer than 200 expressed genes and genes expressed in fewer than 3 cells were filtered out to remove low-quality data. Raw (unnormalized) read counts from both the bulk and single-cell datasets were used as input to the music_prop() function, with free_annotation as the clustering variable and donor as the subject identifier. For ease of interpretation, we then manually grouped the cell types into six biologically meaningful categories: Immune, Endothelial, Muscle, Stromal, and Epithelial.

## Results

### Differential gene expression in muscle tissue

To evaluate the magnitude of sex- and age-specific differences in the transcriptomic profile of the muscle samples of our broader cohort aged 18-80 (N = 67 males aged 45.8 ± 18.8, N = 58 females aged 48.4 ± 19.3), we first visualized variability in gene expression on a PCA plot (Supplementary Fig.1). Ten samples per group (young males aged 18-30, young females aged 18-30 and older males aged 60-75) were then randomly selected for all further analyses along with their corresponding HPMCs.

Forty genes were upregulated, and 46 genes were downregulated in young male when compared to young female muscle tissue (FDR < 0.05, Fig.1A). A Fisher’s exact test indicated that the differentially expressed genes were more likely to be encoded by the Y-chromosome (p < 2.2E-16). Among the most downregulated pathways in males ranked by normalized enrichment scores (NES) were mitochondrial pathway complex I biogenesis, cell replication pathways and several immune pathways. There was no pathway upregulated in males as compared to females (Fig 1B). Unfiltered pathway analysis and corresponding enrichment scores can be found in Supplementary Table 1.

**Figure 1.**
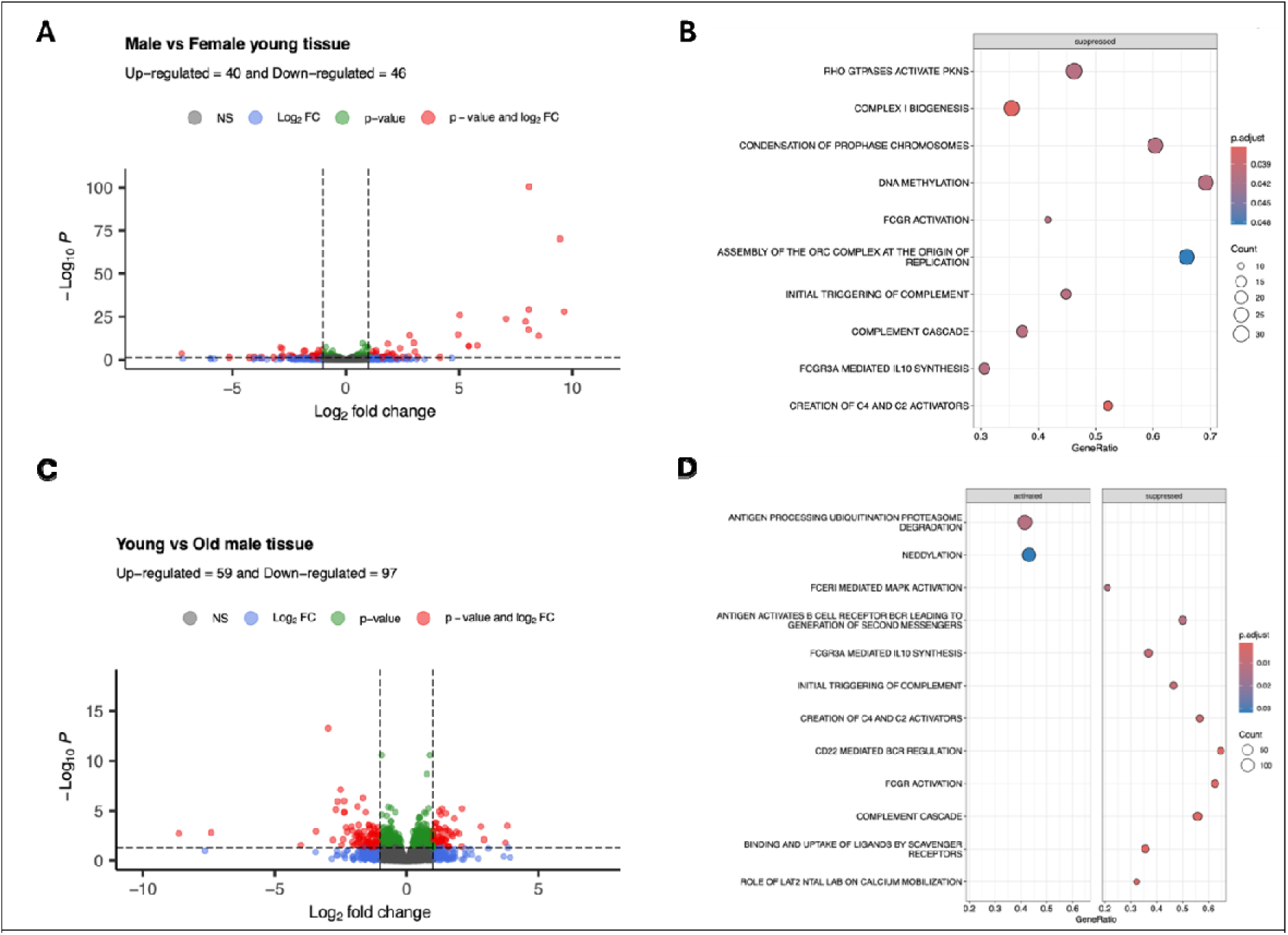
Comparisons between the transcriptomic signature of muscle tissue samples. **A)** Differences in gene expression in young male (control, N=10) and young female (case, N=10) muscle tissue. Genes differentially expressed at FDR < 0.05 and |Log_2_fold change| > 1 are depicted in red. **B)** Reactome pathways upregulated (left panel) or downregulated (right panel) in young male (N=10) *versus* young female (N=10) muscle tissue. **C)** Differences in gene expression in young male (control, N=10) and older male (case, N=10) muscle tissue. Genes differentially expressed at FDR < 0.05 and |Log_2_fold change| > 1 are depicted in red. **D)** Reactome pathways upregulated (left panel) or downregulated (right panel) in young male (N=10) *versus* older male (N=10) muscle tissue.

Fifty-nine genes were upregulated, and 97 genes were downregulated in young male muscle tissue when compared to older male muscle tissue (FDR < 0.05, Fig. 1C). Within the top down-regulated pathways in young males ranked by NES were several pathways related to the immune response and the complement cascade. The two pathways that were upregulated in young *versus* older males were antigen processing ubiquitination proteasome degradation and the related ubiquitin-pathway neddylation (Fig. 1D). Unfiltered pathway analysis and corresponding enrichment scores can be found in Supplementary Table 2.

### Differential gene expression in muscle cell lines

Thirty-two genes were differentially regulated in cell lines grown from young male *versus* young female donors (FDR < 0.05, upregulated: 28, downregulated: 4, Fig. 2A). A Fisher’s exact test indicated that the differentially expressed genes were more likely to be encoded by the Y-chromosome (p < 2.2E-16). Similarly, nine genes were differentially regulated in cell lines grown from young *versus* older male donors (FDR < 0.05, upregulated: 3, downregulated: 6, Fig. 2B). In both cases, gene set enrichment analysis (GSEA) resulted in no significantly enriched pathways.

**Figure 2.**
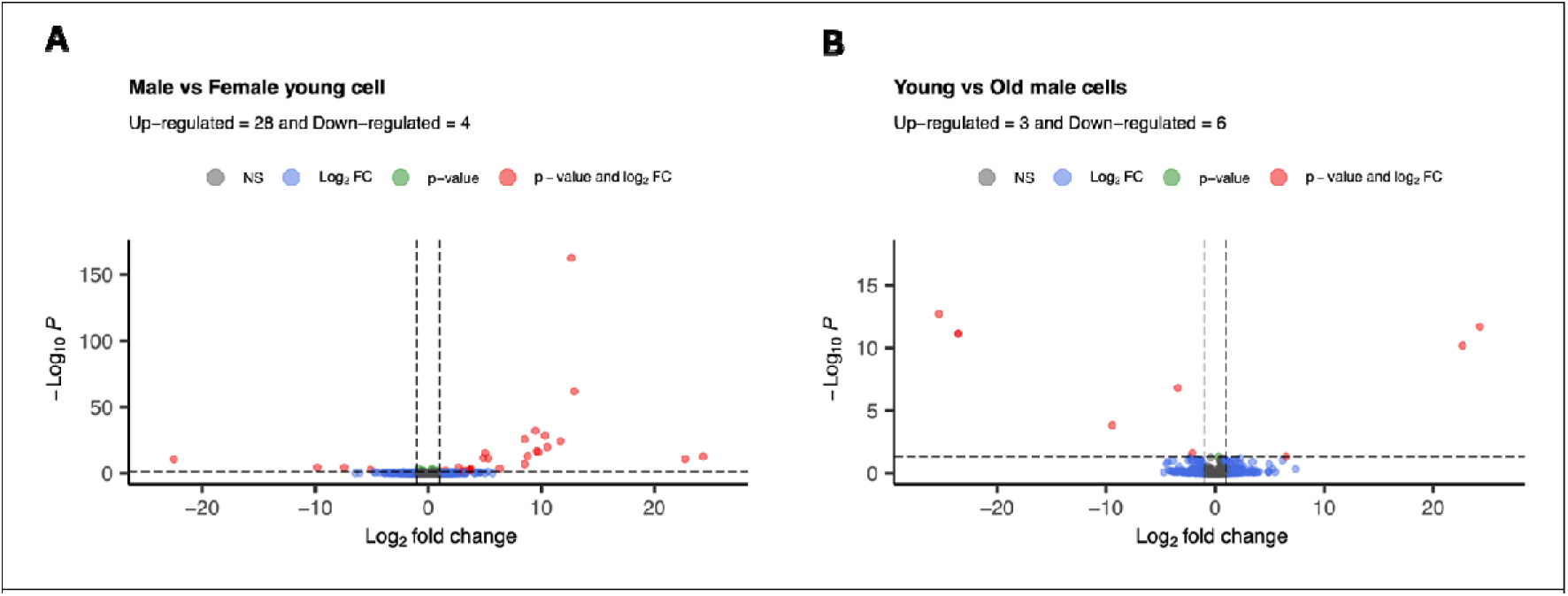
Comparisons between the transcriptomic signature of HPMCs. **A)** Differences in gene expression in HPMC lines grown from young male (control, N = 10) *versus* young female (case, N = 10) donors. **B)** Differences in gene expression in HPMC lines grown from young male (control, N = 10) *versus* older male (case, N = 10) donors. Genes differentially expressed at FDR < 0.05 and |Log_2_fold change| > 1 are depicted in red.

### Comparison of the transcriptome of muscle tissue and HPMC lines

Differences in the transcriptome of our subset of 30 tissue samples *versus* their corresponding cell lines were first visualized using principal component analysis (PCA). It revealed a striking separation by sample type (tissue or HMPCs) along the first principal component (88.6%) (Fig. 3A), which was explained by 9595 genes being differentially regulated between muscle tissue samples and their corresponding HPMC lines (Fig. 3B). Tissue samples formed a tight cluster, while HPMC lines demonstrated a broader spread along the first and second principal component. By restricting the PCA to HPMC only, male and female cell lines separated by sex along the second principal component (Supplementary Fig. 2A). Similarly, performing PCA solely on tissue samples revealed sex-specific separation along the first and second principal components (Supplementary Fig. 2B).

**Figure 3.**
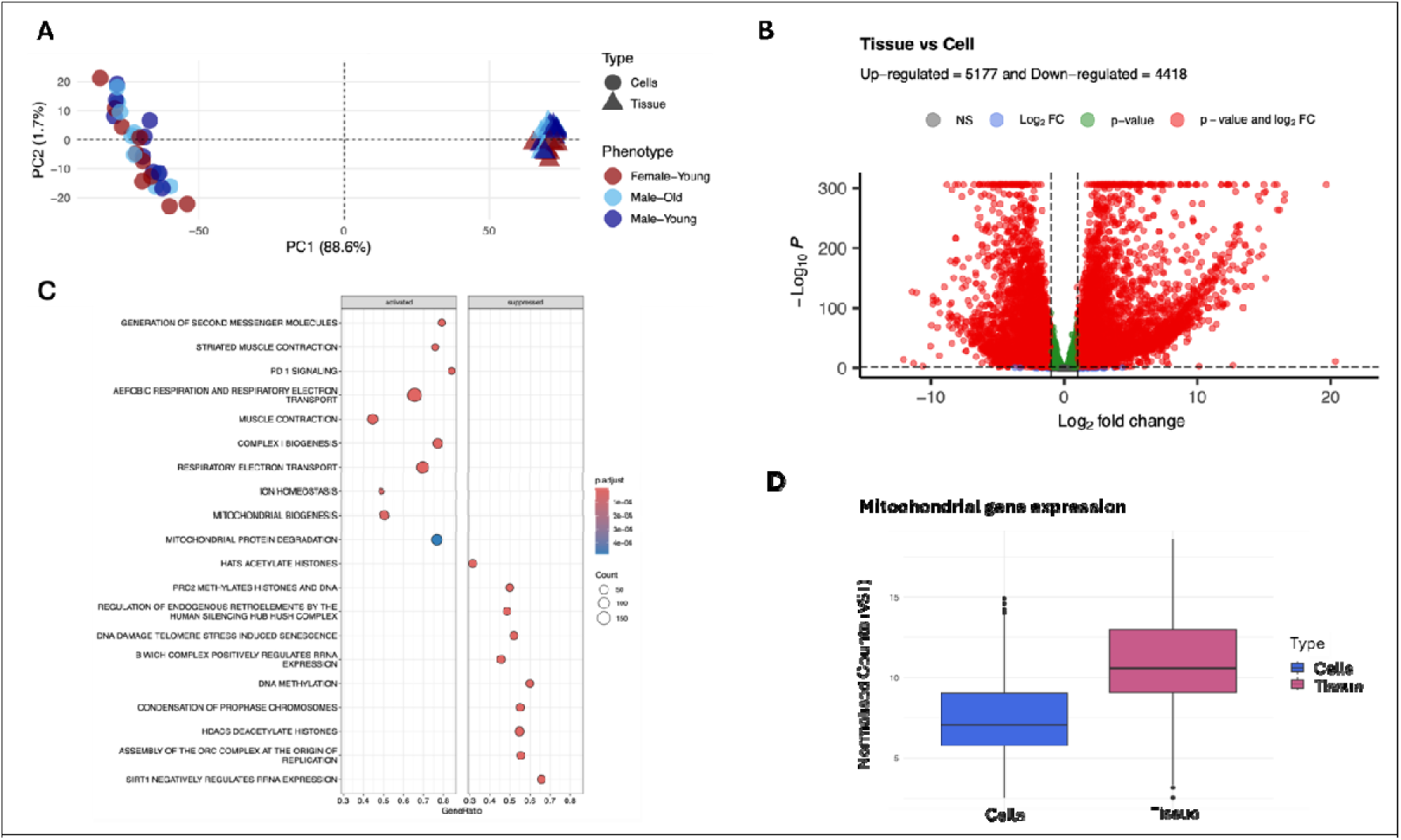
Comparisons between the transcriptomic signature of muscle tissue and HPMCs. **A)** Principal component analysis of the transcriptome of N = 30 human muscle tissue samples (control, N = 10 young females, N = 10 young males, N = 10 older males) and the corresponding N = 30 HPMC lines grown from the same donors (case). **B)** Differences in gene expression in human muscle tissue (N=30) compared to primary muscle cell lines grown from the same donors (N=30). Genes differentially expressed at FDR < 0.05 and |Log_2_fold change| > 1 are depicted in red. **C)** Reactome pathways upregulated (left panel) or downregulated (right panel) in muscle tissue (N=30) *versus* HPMCs (N=30). **D)** Average sum of normalised counts of the 13 mitochondrial protein-coding genes in cell (blue) and tissue (red) samples.

The main pathways differentially expressed between HPMCs and muscle tissue pertained to energy metabolism (e.g. aerobic respiration and respiratory electron transport, complex I biogenesis, mitochondrial biogenesis and transcriptional activation of mitochondrial biogenesis were upregulated in tissue *versus* cells); muscle contraction and striated muscle contraction (upregulated in tissue *versus* cells); and, interestingly, epigenetic modifications (e.g. DNA methylation, HDACs deacetylate histones, HATS acetylate histones and negative epigenetic regulation of rRNA expression were downregulated in tissue *versus* cells) (Fig. 3C). Unfiltered pathway analysis and corresponding enrichment scores can be found in Supplementary Table 3. The enrichment of mitochondrial pathways could be primarily attributed to an increase in expression of all 13 mitochondrial-encoded genes in tissue *versus* cells (Fig. 4D and Supplementary Fig. 3).

**Figure 4.**
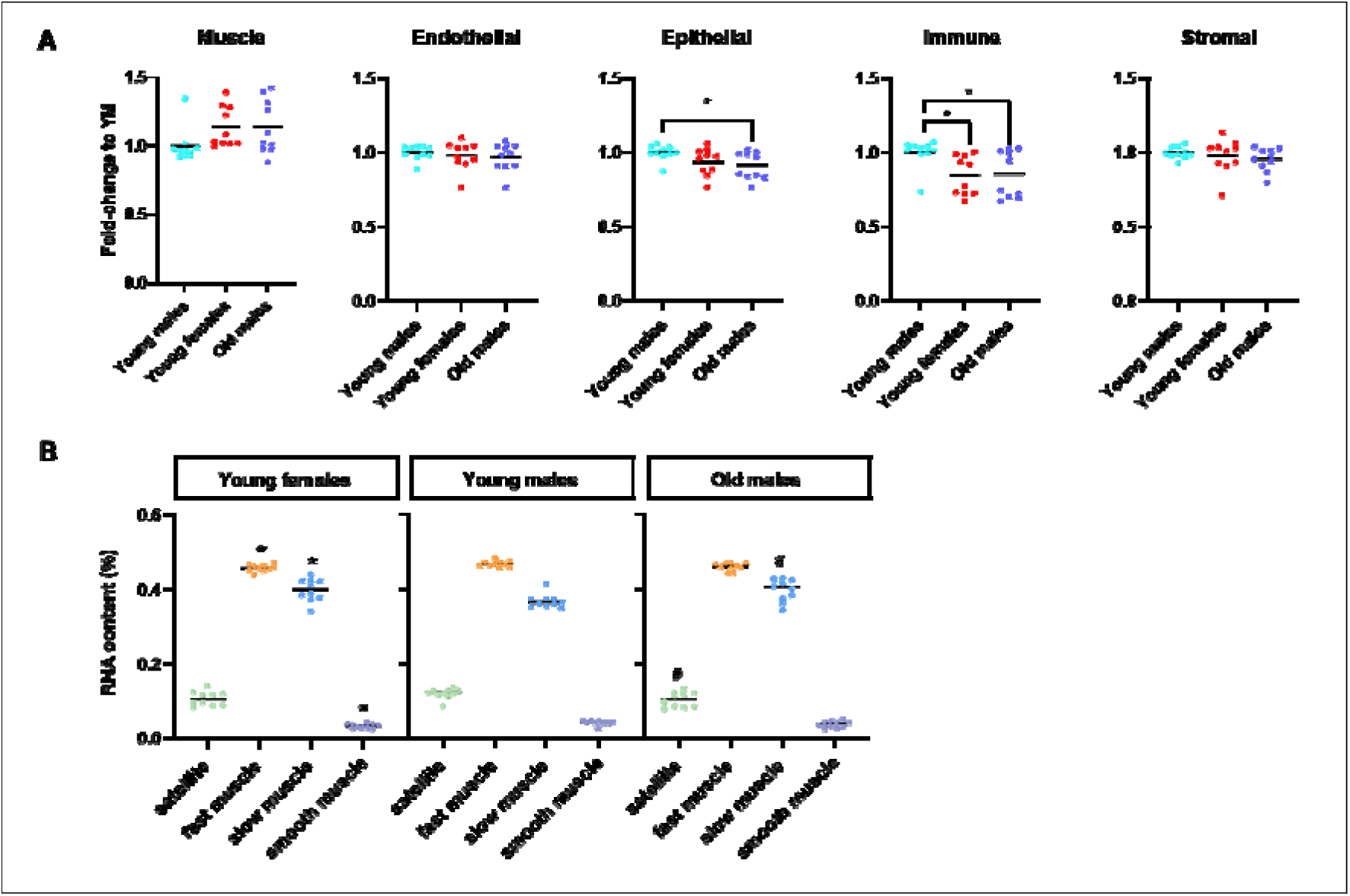
Deconvolution of bulk-RNASeq results in muscle tissue. **A)** Relative amount of muscle, endothelial, epithelial, immune and stromal cell RNA when compared to the young male group. * denotes p < 0.05. **B)** Proportion of RNA from satellite, fast, slow and smooth muscle cell origin in each sample group. * and # denote p < 0.05 when compared to the young male group.

### Cell type deconvolution in muscle tissue

To understand the potential influence of non-myogenic resident cell populations (28, 29) on our bulk RNASeq outputs, we next deconvoluted our muscle tissue transcriptome results using Multi-subject Single Cell deconvolution (MuSIC). Due to the unique multinucleated nature of skeletal muscle cells, there is currently no method allowing quantitative deconvolution of bulk RNA-Seq data in skeletal muscle (30, 55), however, MuSIC and others algorithms enable robust semi-quantitative comparisons between groups of samples, or between different muscle sample populations (30).

Muscle tissue samples comprised a mix of RNA from muscle (including fast, slow, smooth and satellite muscle cell), endothelial (including capillary, arterial and venous endothelial cells), immune (including t-cells, natural-killer cells, mast cells, macrophages and neutrophils), stromal (including mesenchymal cells and pericytes) and epithelial cells (Figure 4A). There was no difference in the amount of muscle RNA content between any group. Subtle differences in the relative proportion of non-myogenic, mononucleated cell populations however existed between groups, with an 8.8% decrease in the amount of RNA from epithelial cells in older *versus* young muscle tissue, and a 15% decrease in the amount of RNA from immune cells in both older *versus* young muscle tissue and young female *versus* young muscle tissue (all p < 0.05). Within muscle cell populations, fast muscle RNA was most prominent (46% of all muscle RNA), followed by slow muscle RNA (39%) and RNA from satellite cells (11%) (Figure 4B). As expected, there was significantly more fast muscle RNA in young male than in young female tissue, and significant less slow muscle RNA in young male than in both young female and older male female tissue. Finally, there was less satellite cell RNA in older than in young muscle tissue (all p<0.05).

### Transcriptomic signature of myocyte differentiation

Harvesting thirty cell lines at the same time point may generate some degree of variability in the differentiation level of individual cell lines. Because the degree of satellite cell differentiation into myotubes is complex to assess objectively, we used an existing RNA-seq dataset generated by our group to establish a differentiation signature of human primary myocytes. Rather than relying on a single measure (e.g. fusion index) or a limited number of muscle differentiation markers (e.g. MyoD, MyoG1), we sought to assess the individual level of differentiation of each HPMC line at our experimental endpoint when compared to two reference cell lines. This was done by comparing each cell line transcriptomic profile to the dynamic signature extracted from an existing, tightly controlled time-course experiment conducted under the same conditions. To define a reference transcriptomic differentiation signature, we first identified genes that were differentially expressed (FDR<0.05) at a given timepoint from the onset of differentiation (T2) comparing each time point to the next (T2 vs T3, T3 vs T4, T4 vs T5, T5 vs T6, Fig. 5A). This analysis resulted in 1411 unique genes (or 8% of all detected genes) that could be grouped in 14 clusters containing 29-773 genes (Fig. 5B). Associated Reactome pathways for the first 10 clusters can be found in Supplementary Fig. 4. The top 3-clusters were respectively enriched in genes related to the cell cycle (cluster 1), lipid metabolism (cluster 2) and muscle contraction (cluster 3).

**Figure 5.**
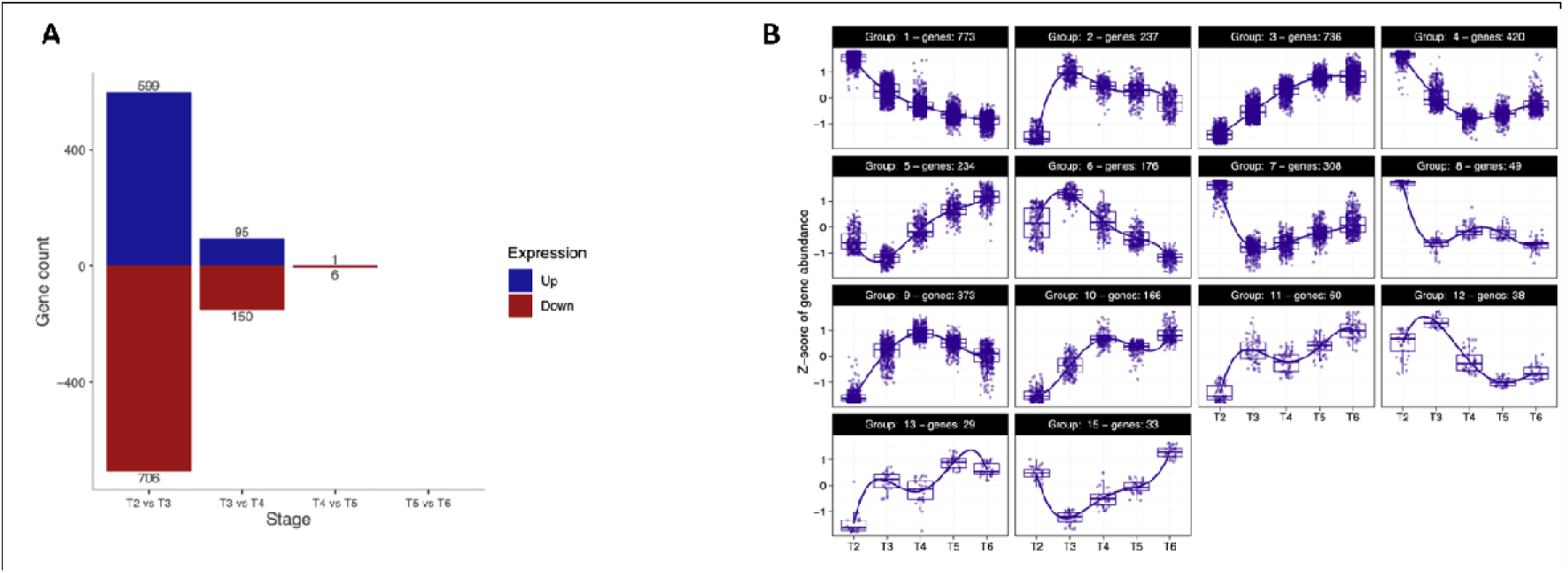
Differentiation time course. **A)** Genes differentially expressed between each differentiation time point and the following one (P < 0.05). **B)** Expression profiles of each gene differentially regulated between each differentiation time point and the following one were grouped into 15 clusters containing 29-773 genes.

The transcriptomic profile of our differentiation time course served as a basis to generate a differentiation index comprised between 0-1, where the least differentiated cell lines of the time course were assigned score closer to 0 and the most differentiated cell lines of the time course were assigned a score closer to 1 (Supplementary Fig. 5). This index was then applied to each HPMC line. Differentiation scores (Supplementary Table 4) for each cell line were comprised between 0.96 and 1.09 (mean ± SD = 1.03 ± 0.04).

To test whether differences in the differentiation status of individual cell lines explained any of the variability, we conducted all further analyses with and without individual differentiation indices as a co-variate in all our models. Results remained the same for all analyses, and therefore, the results presented below are not adjusted for differentiation status.

### Maintenance of the age signature in HPMC lines from male donors

Each individual gene expressed in young and older muscle tissue and in young and older muscle cell lines was plotted on a scatter plot regardless of p-value (Supplementary Fig. 6A). Genes whose differential expression failed to reach statistical significance (FDR < 0.05, log2FC > 1, < -1) between young and older muscle tissue and/or young and older muscle cell lines (represented in yellow in Supplementary Fig. 6A) were then subtracted from the plot (Fig. 6A). Overall, the overlap in differentially expressed genes between tissues and cell lines was minimal. Of the remaining genes, only one gene, *HLA-C*, was significantly downregulated in young when compared to older tissue, as well as in young when compared to older cell lines. This was reflected by a Spearman’s correlation coefficient of 0.001 from the DESeq2 test statistic of all genes, denoting a lack of association between the two transcriptomic profiles. This lack of association can be visualized on the corresponding contour plot (Fig. 6B) that shows genes spread relatively evenly across the four quadrants. Notably, the density contours suggest the absence of a strong directional bias, with no quadrant showing dominant clustering of genes. A modest concentration of genes appears in the bottom-right quadrant, indicating a subset of genes that are upregulated in the “Cell” contrast but downregulated in the “Tissue” contrast. However, this pattern is not pronounced, and the overall dispersion indicates that gene expression changes associated with aging in cultured muscle cells do not systematically mirror those observed in muscle tissue.

**Figure 6.**
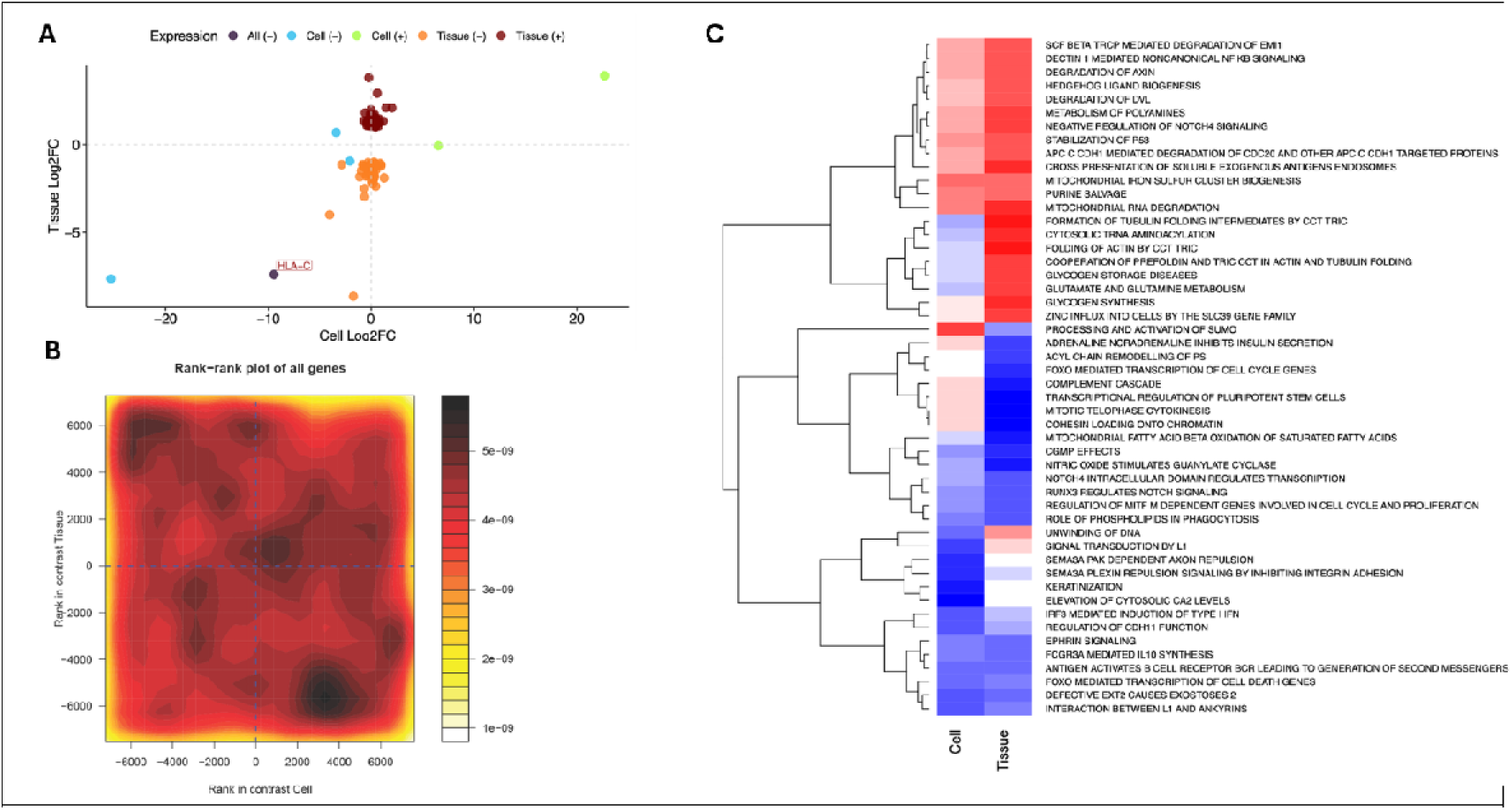
Maintenance of the age phenotype between muscle tissue and HPMCs. **A)** Scatter plot depicting the log_2_fold-change of each individual gene expressed in young (N=10) and older (N=10) muscle tissue and in the corresponding young (N=10) and older (N=10) muscle cell lines. Genes differentially expressed at FDR < 0.05 and down regulated in both young cell and muscle tissue are depicted in black. Genes differentially expressed at FDR < 0.05 and down regulated in young cells only are depicted in blue. Genes differentially expressed at FDR < 0.05 and up regulated in young cells only are depicted in green. Genes differentially expressed at FDR < 0.05 and down regulated in young muscle tissue only are depicted in orange. Genes differentially expressed at FDR < 0.05 and up regulated in young muscle tissue only are depicted in dark red. **B)** Differential expression rank–rank density contour plot with cell and tissue contrasts on the x- and y-axes, respectively. Each point represents a gene ranked by its differential expression in both contrasts, and the filled contours indicate gene density across the rank space. The four quadrants correspond to directional patterns of regulation: genes upregulated in both contrasts (top right), downregulated in both (bottom left), or oppositely regulated (top left and bottom right). The relatively even spread of density across all quadrants reflects the absence of a consistent transcriptional signature shared between aged muscle tissue and cultured cells. **C)** Heatmap depicting the differentially regulated Reactome pathways in young (N=10) and older (N=10) muscle tissue (control) and in the corresponding young (N=10) and older (N=10) HPMC lines (case).

A similar analysis was then repeated at the pathway level. We used *mitch* to compare the biological pathways that were significantly regulated in the same or in the opposite direction between young and older muscle tissue and/or the corresponding young and older HPMC lines. The top-50 significant pathways classified by s-distance were visualized on a heatmap (Fig. 6C, see Supplementary Table 5 for the complete, non-filtered list of pathways). When comparing muscle tissue and HPMC lines, 31 pathways were regulated in the same direction with ageing, and 19 pathways were regulated in the opposite direction with ageing. Differentially regulated pathways included those involved in the regulation of structural proteins, glycogen synthesis and storage, and cell proliferation.

### Maintenance of the sex signature in HPMC lines from young donors

Similarly, each gene expressed in young male and female muscle tissue and in young male and female cell lines was plotted on a scatter plot regardless of significance (Supplementary Fig. 6B), including genes encoded on the sex chromosomes. Genes whose differential expression failed to reach statistical significance (FDR < 0.05, log2FC > 1, < -1) between young male and female muscle tissue and/or young male and female muscle cell lines (represented in orange in Supplementary Fig. 6B) were then subtracted from the plot (Fig. 7A). Two genes, *HLA-C* and *FEZF2*, were significantly downregulated in male tissue when compared to female tissue, as well as in male HPMC when compared to female HPMC and 11 genes (*TTTY14, PRKY, USP9Y, NLGN4Y, TXLNGY, KDM5D, ZFY, UTY, E1F1AY, DOX3Y, RPS4Y1*) were significantly upregulated in male tissue when compared to female tissue, as well as in male HPMCs when compared to female HPMCs. Spearman’s correlation coefficient was -0.06, denoting a weak, negative association of the test statistics between the two transcriptomic profiles, which can be visualized on the corresponding contour plot (Fig. 7B). The density contours show a modest concentration of genes in the bottom-right and top-left quadrants, indicating a subset of genes that are upregulated in cells but downregulated in tissue, and vice versa. In contrast to the age comparison, the sex-associated expression differences displayed a more evident pattern of discordant regulation between in vivo and in vitro models. These opposing trends are consistent with the weak negative correlation observed and suggest that sex-biased expression is not preserved across tissue and cell models. These figures capture distinct but complementary dimensions of the data, which revealed a positive correlation between the log-fold change of a few genes while some of the remaining genes exhibited a negative association.

**Figure 7.**
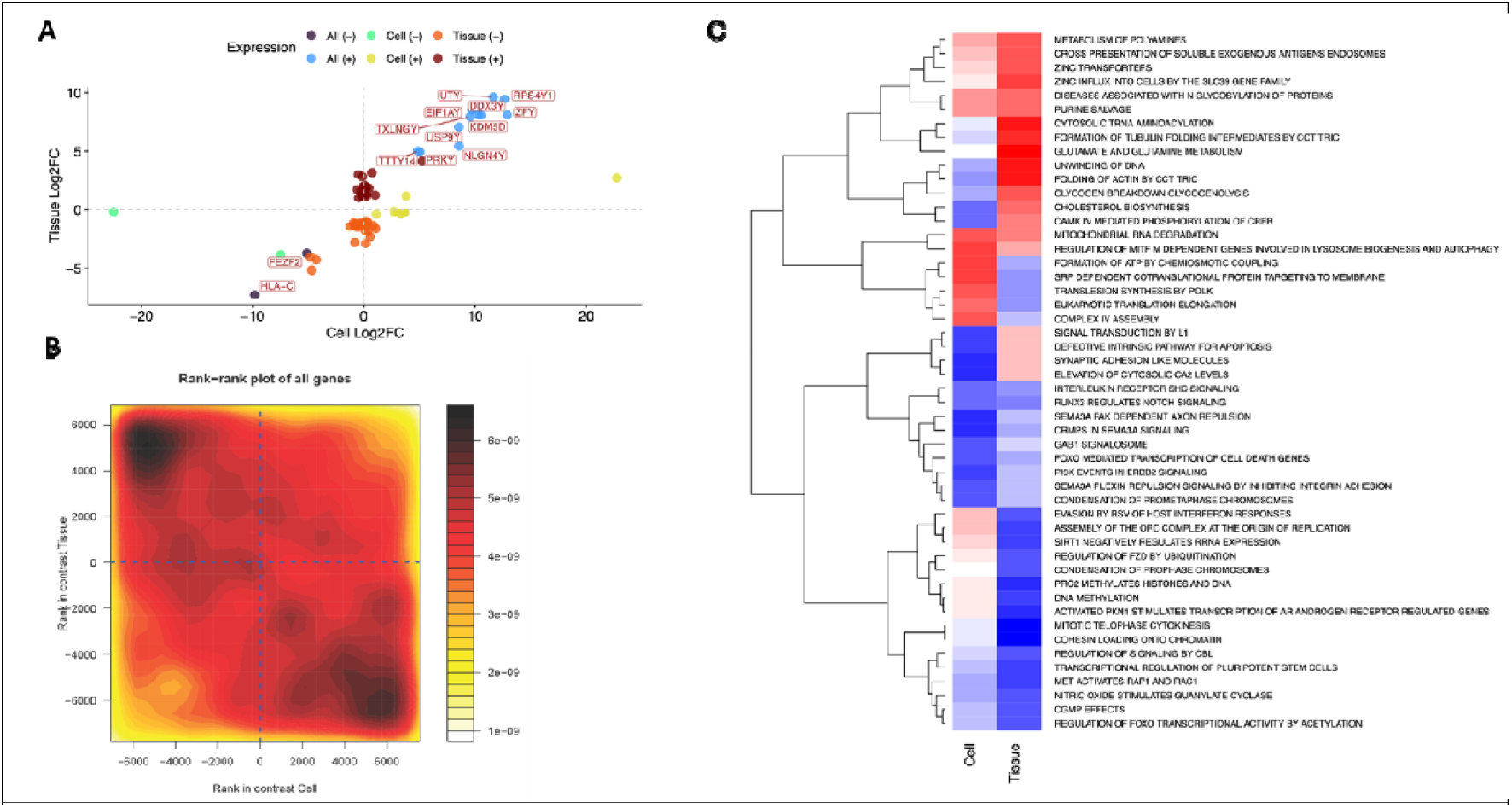
Maintenance of the sex phenotype between muscle tissue and HPMCs. Scatter plots depicting the log_2_fold-change of each individual gene expressed in young male (N=10) and young female (N=10) muscle tissue and in the corresponding young male (N=10) and young female (N=10) muscle cell lines. Genes differentially expressed at FDR < 0.05 and down regulated in both female cell and muscle tissue are depicted in black. Genes differentially expressed at FDR < 0.05 and upregulated in both male cell and male muscle tissue are depicted in blue. Genes differentially expressed at FDR < 0.05 and down regulated in male cells only are depicted in aqua. Genes differentially expressed at FDR < 0.05 and down regulated in male muscle tissue only are depicted in dark orange. Genes differentially expressed at FDR < 0.05 and up regulated in male muscle tissue only are depicted in dark red. **B)** Differential expression rank–rank density contour plot with cell and tissue contrasts on the x- and y-axes, respectively, as described in Fig. 6B. In this case, gene density is enriched in the top-left and bottom-right quadrants, indicating opposing patterns of sex-biased gene expression between tissue and cell models. **C)** Heatmap depicting the differentially regulated pathways in young male (N=10) and female (N=10) muscle tissue (control) and in the corresponding young male (N=10) and female (N=10) muscle cell lines (case).

A similar analysis was then repeated at the pathway level. The top-50 significant pathways classified by s-distance were visualized on a heatmap (Fig. 7C, see Supplementary Table 6 for the complete, non-filtered list of pathways). Amongst those, 25 pathways were regulated in the same direction with sex in both tissues and HPMC, and 25 pathways were regulated in opposite directions.

## Discussion

First isolated in 1981 (2), HPMCs constitute a common experimental model in muscle research due to the relative ease of access to human skeletal muscle tissue using well-established percutaneous biopsy techniques (56). Despite their intrinsic genetic heterogeneity, HPMCs are often preferred to the commercially available mouse C_2_C_12_ or rat H6 muscle cell lines due to their non-immortalized nature and perceived closeness to a human physiological system. Indeed, there are important transcriptional and metabolic differences between HPMCs, C_2_C_12_ and H6 myocytes (5). However, the variability between the transcriptome of muscle tissue and cultured myocytes from the same species (human, mouse or rat) is substantially larger than the variability between the muscle tissue transcriptomes from different species, or between the transcriptome of cultured myocytes from different species (5). An extension of the premise that HPMCs accurately mimic a human physiological system is the common use of HPMC lines that share traits with the population of interest (e.g. “aged” muscle cell lines, “female” muscle cell lines) to probe age- or sex-specific mechanisms. The assumption underlying such practise is that HPMCs may retain some key phenotypical characteristics from their donors once placed in culture despite the removal of the systemic environment. Here we directly tested this assumption by conducting an in-depth comparison of the transcriptome of young male, young female, and older male muscle tissue to their corresponding HPMCs at the individual gene and pathway level.

In line with what was previously reported by us and others, we found differences between the transcriptome of young male *versus* female muscle tissue (6–10), as well as between young *versus* older male muscle tissue (21–24). The discovery of sex-specific differences during the ageing process was beyond the scope of this study. Despite differences between studies pertaining to the type of muscle used, the size of the cohort, the time of sample collection and the age of the participants, and the stringency of the statistical threshold applied, sex- and age-specific pathway analysis revealed strong overlap between our results and the existing literature. For example, our results recapitulated those of previous studies having reported age differences in the immune response (21, 57) and dysregulation of the ubiquitin proteasome complex (58). Similarly, sex-differences in muscle mitochondrial activity (7, 23) are well described in the literature.

Generally, these differences did not translate to the corresponding HPMC lines. Thirty-two individual genes were differentially regulated in HPMCs grown from young male *versus* young female donors (confirming micro-array based findings from (27)) and nine individual genes were differentially regulated in HPMCs grown from young *versus* older male donors, respectively. We tested the maintenance of the age and sex phenotype by investigating how many genes remained differentially regulated in the same direction between two groups of tissue samples (e.g. young *versus* older males) and their corresponding two groups of cell lines. Regarding age, only one gene (*HLA-C*) remained differentially regulated in the same direction in both muscle tissue and HPMCs (Spearman’s correlation coefficient = 0.001), with *HLA-C* gene expression downregulated by > 5-fold in tissue in young *versus* older tissue samples, and by > 10-fold in young in *versus* older cell samples. Regarding the maintenance of the sex phenotype, 13 genes (*TTTY14, PRKY, USP9Y, NLGN4Y, TXLNGY, 4DM50, ZFY, UTY, E1F1AY, DOX3Y, RPS4Y1, FEZF2, HLA-C*) remained differentially regulated in the same direction in both tissue and HPMCs, despite a negative Spearman’s correlation coefficient (-0.06). This strongly suggests that the expression of some other significantly regulated genes, which were not specifically investigated in this study, must have exhibited a reversed association between tissue and their corresponding HPMCs; a hypothesis supported by our pathway analysis.

*HLA-C,* the only gene whose differential expression was maintained with age, encodes a protein belonging to the major histocompatibility complex (MHC) class I family and was significantly downregulated in young *versus* older samples in both tissue and cells. An altered immune response (also termed “immunosenescence”) is both a consequence and an indirect hallmark of ageing (59), including at the transcriptomic level (22). Ageing is however also characterized by a constitutive, low-level activation of the immune system (60), also detectable at the proteomic level (57), which may explain that some immune features increase rather than decrease with age. Pathway analysis offers higher-level insights into the maintenance of the muscle immune phenotype with ageing in tissue and HPMCs. While HLA-C was regulated in the same direction in tissue and HPMCs, the complement cascade, for example, was downregulated in young *versus* older tissue, but upregulated in young *versus* older HPMCs. These disparities may be due to the fact that whole skeletal muscle tissue inevitably contains resident immune cells (29). Indeed, we found RNA originating from T-cells, natural-killer cells, mast cells, macrophages and neutrophils in our muscle tissue sample bulk RNA, in amounts that significantly differed between groups. These cells are expected to be filtered out by the muscle specific anti-CD56+ antibodies used in the process of establishing HMPCs, but, in tissue samples, may influence individual gene expression across certain cellular pathways such as those related to immunity. However, other striking contrasts that may not be directly attributed to differences in cell populations were also evident in the cellular pathways pertaining to the regulation of structural muscle proteins, or glycogen metabolism.

All 11 genes that remained up-regulated in both male *versus* female tissue and cells were Y-encoded. Of the two genes that remained down-regulated in both male *versus* female tissue and cells, FEZF2 is a sexually dimorphic gene encoding a transcription factor important to cell fate in the development of the nervous system in mammals (61, 62). In turtle gonads, which are subjected to temperature-dependent sex-determination, FEZF2 is hypermethylated in females *versus* males (63), suggesting a role in sexual differentiation, but how this may translate to other tissues and species remains unknown.

These broad transcriptomic differences between human muscle tissue and their corresponding HMPCs may be explained in two ways. Firstly, in mammals, skeletal muscle fibres make up to 95% of the volume of muscle tissue but display relatively sparse nuclei when compared to their cell size. In addition, whole muscle biopsies not only contain myoblasts and mature muscle cells, but also fibro-adipogenic progenitors, immune cells, fibroblasts and endothelial cells (28, 29), which was consistent with our bulk RNA deconvolution results. Early studies have suggested that proportion of muscle nuclei may be as little as 50% (64, 65) even though myocytes remain by far the most transcriptionally active cell type in the muscle (30, 66). Whether potential differences in cell type proportions between our muscle sample groups may have accounted for some of the variability was investigated using a deconvolution algorithm. Deconvolution methods applicable to bulk RNASeq are relatively new (55) and have not been optimized for skeletal muscle tissue (30). Indeed, muscle fibres are large, multinucleated syncytia that contribute a disproportionate amount of RNA when compared to mono-nucleated cells, which can mask the signal from smaller or less transcriptionally active cell types like satellite cells, immune cells, or endothelial cells. These methods can however provide accurate semi-quantitative comparisons within a same cell group or between different sample groups (30). Here, we found no change in muscle RNA content between any of our groups, and some subtle but significant differences in the relative proportions of epithelial and immune cells, which may have had marginal effects on our between-group comparisons. These aspects must therefore be considered when comparing patterns of gene expression between whole muscle tissue and cultured muscle cells that have been subjected to muscle-specific cell selection.

Secondly, the hormonal environment drastically varies between young males and females, exerting a sex-specific effect on numerous tissues including skeletal muscle (reviewed in (67)). Circulating concentrations of androgenic and oestrogenic hormones are highly sex specific. In addition, these hormones enact their effects through specific receptors that can be differentially regulated with sex (68) or show sex-biased activation patterns (9). Our results indicate that the removal of the sex-determining hormonal milieu is enough to conceal most transcriptomic differences due to sex *in vitro*, including some of the differences mediated by the Y chromosome complement independent of sex hormones (26, 69). The same may be true, to an extent, with the ageing process. In males, ageing is associated with profound declines in androgen concentrations, particularly testosterone (70, 71). The direct effects of testosterone on age-related molecular mechanisms in the muscle are yet to be determined, especially *in vivo*, but evidence suggests that testosterone decline impacts several pathways important to the maintenance of skeletal muscle cells with ageing. These include muscle fibre hypertrophy through the suppression of ubiquitin-ligase mediated atrophic pathways, or satellite cell proliferation through the synthesis of cell-cycle regulating proteins such as follistatin and Notch receptors (72, 73). Prior research in C_2_C_12_ has mimicked aspects of ageing in cells, where subjecting the cells to multiple-population-doubling resulted in impaired myoblast fusion and upregulated myostatin and TNF-α (74); effects that could be partially rescued by testosterone treatment (75, 76). Multiple-population-doubling is however not applicable to HPMCs, and the lack of androgens and other growth factors in the cellular milieu suggests a major role for these hormones in the determination of the ageing muscle phenotype. Conversely, the presence of foetal growth factors in the proliferation medium may push the cells towards a “younger” phenotype.

In conclusion, we provide evidence that the transcriptomic signature of sex and age in human muscle tissue is mostly lost once cultured *in vitro*. At the individual gene level, this loss of phenotype is as prominent for age as it is for sex, despite the maintained differential expression of Y-linked genes, whose sex-specific expression was more likely to remain conserved in both tissue and cell lines. The transcriptome only provides insights into one layer of metabolic and functional regulation though, and these results should be further validated, for example at the proteomics, phenotypic and function levels, before discounting the use of HMPCs as age- or sex-specific models of human skeletal muscle. Our findings also do not exclude the possibility that some sex- or age-specific phenotypes might be lost at baseline but become apparent in response to an external stimulus (e.g. hormone treatment, electrostimulation, miRNA transfection) as previously suggested by our group (77). It is also possible that cell culture media may be supplemented with specific age- and sex-specific circulating growth factors to achieve a model transcriptionally closer to the source phenotype. Finally, nuclear single cell sequencing or HMPCs grown in 3D culture as organoids under strain might provide a closer comparison between human muscle fibres and their corresponding HPMCs. Our results generally call for caution, and any claim that HPMCs can be used as an experimental model of human muscle sex or age should be interpreted carefully.

## Declarations

### Ethics approval and consent to participate

This research study was granted ethical approval by the Deakin University Human Research Ethics Committee (DUHREC 2021-307).

### Consent for publication

N/A

### Data availability statement

The RNA-seq data generated and analysed in this study have been deposited in the Gene Expression Omnibus (GEO) under accession number GSE287342. The full code is available at https://github.com/megan-soria/age_and_sex_signature. The *contam* package is available at https://github.com/markziemann/contam/tree/main.

### Authors’ contributions

SL designed the study, analysed the data, interpreted the data and funded the study. MS analysed the data and interpreted the data. RW, AC, KVB and AG collected the samples and generated the data. AV and TB generated and analysed the differentiation data. DH collected the samples, analysed the data and interpreted the data. MZ designed the study and interpreted the data. All authors contributed to the preparation of the manuscript.

## Acknowledgements

The authors would like to acknowledge Dr Sarah Voisin (Novo Nordisk) for her precious advice. This research was supported by use of the Nectar Research Cloud, a collaborative Australian research platform supported by the NCRIS-funded Australian Research Data Commons (ARDC). The authors gratefully acknowledge the contribution to this work of the Victorian Operational Infrastructure Support Program received by the Burnet Institute.

## Figure Legends

Supplementary Figure 1. Principal component analysis of the transcriptome of N = 125 human muscle tissue samples from participants aged 18-80 (N = 58 females, N = 67 males). Female are represented in red and male are represented in blue. Increased colour opacity indicates increased age.

Supplementary Figure 2**. A)** Principal component analysis of the transcriptome of N = 30 HPMC lines. Female are represented in red and male are represented in blue. Young donors (18–30) are represented by circles and older donors (60–75) are represented by triangles. **B)** Principal component analysis of the transcriptome of N = 30 muscle tissue samples. Female are represented in red and male are represented in blue. Young participants (18–30) are represented by circles and older participants (60–75) are represented by triangles.

Supplementary Figure 3. Average sum of normalised counts of mitochondrial protein-coding genes MT-ND2, MT-ND3, MT-ND4, MT-ND5, MT-ND6, MT-ND4L, MT-ATP8, MT-CO1, MT-CO2, MT-CO3, MT-CYB, MT-ND1 and MT-ATP6 in cell (blue) and tissue (red) samples.

Supplementary Figure 4. Pathway analysis of the top-10 clusters identified across the differentiation time course.

Supplementary Figure 5. Differentiation scores across timepoints according to weighted sum (PCA1). The earlier differentiation time points are assigned a score closer to 0 and the later differentiation time points are assigned a score closer to 1.

Supplementary Figure 6**. A)** Scatter plot depicting the log_2_fold-change of each individual gene expressed in young (N=10) and older (N=10) muscle tissue and in the corresponding young (N=10) and older (N=10) muscle cell lines. Genes differentially expressed at FDR < 0.05 and down regulated in both young cell and muscle tissue are depicted in black. Genes differentially expressed at FDR < 0.05 and down regulated in young cells only are depicted in blue. Genes differentially expressed at FDR < 0.05 and up regulated in young cells only are depicted in green. Genes differentially expressed at FDR < 0.05 and down regulated in young muscle tissue only are depicted in orange. Genes differentially expressed at FDR < 0.05 and up regulated in young muscle tissue only are depicted in dark red. Genes not differentially expressed between cells and tissues are depicted in yellow.

Supplementary Figure 6**. B)** Scatter plots depicting the log_2_fold-change of each individual gene expressed in young male (N=10) and young female (N=10) muscle tissue and in the corresponding young male (N=10) and young female (N=10) muscle cell lines. Genes differentially expressed at FDR < 0.05 and down regulated in both female cell and muscle tissue are depicted in black. Genes differentially expressed at FDR < 0.05 and upregulated in both male cell and male muscle tissue are depicted in blue. Genes differentially expressed at FDR < 0.05 and down regulated in male cells only are depicted in aqua. Genes differentially expressed at FDR < 0.05 and down regulated in male muscle tissue only are depicted in dark orange. Genes differentially expressed at FDR < 0.05 and up regulated in male muscle tissue only are depicted in dark red. Genes not differentially expressed between cells and tissues are depicted in orange.

