## Supplementary Figures for "The transcriptomic signature of age and sex is not conserved in human primary myocytes"

### Slide 1
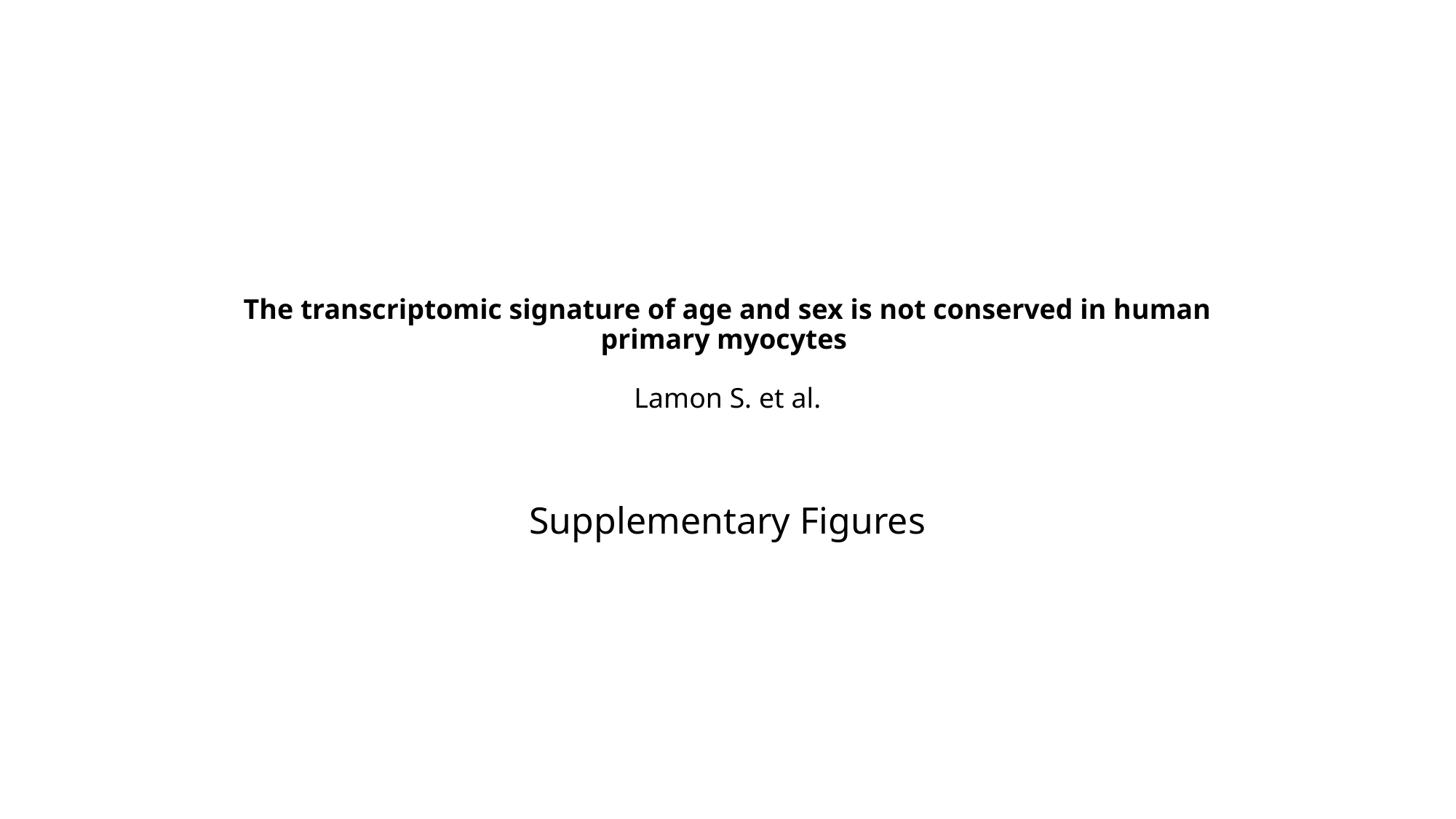

#### The transcriptomic signature of age and sex is not conserved in human primary myocytes  Lamon S. et al.
Supplementary Figures

### Slide 2
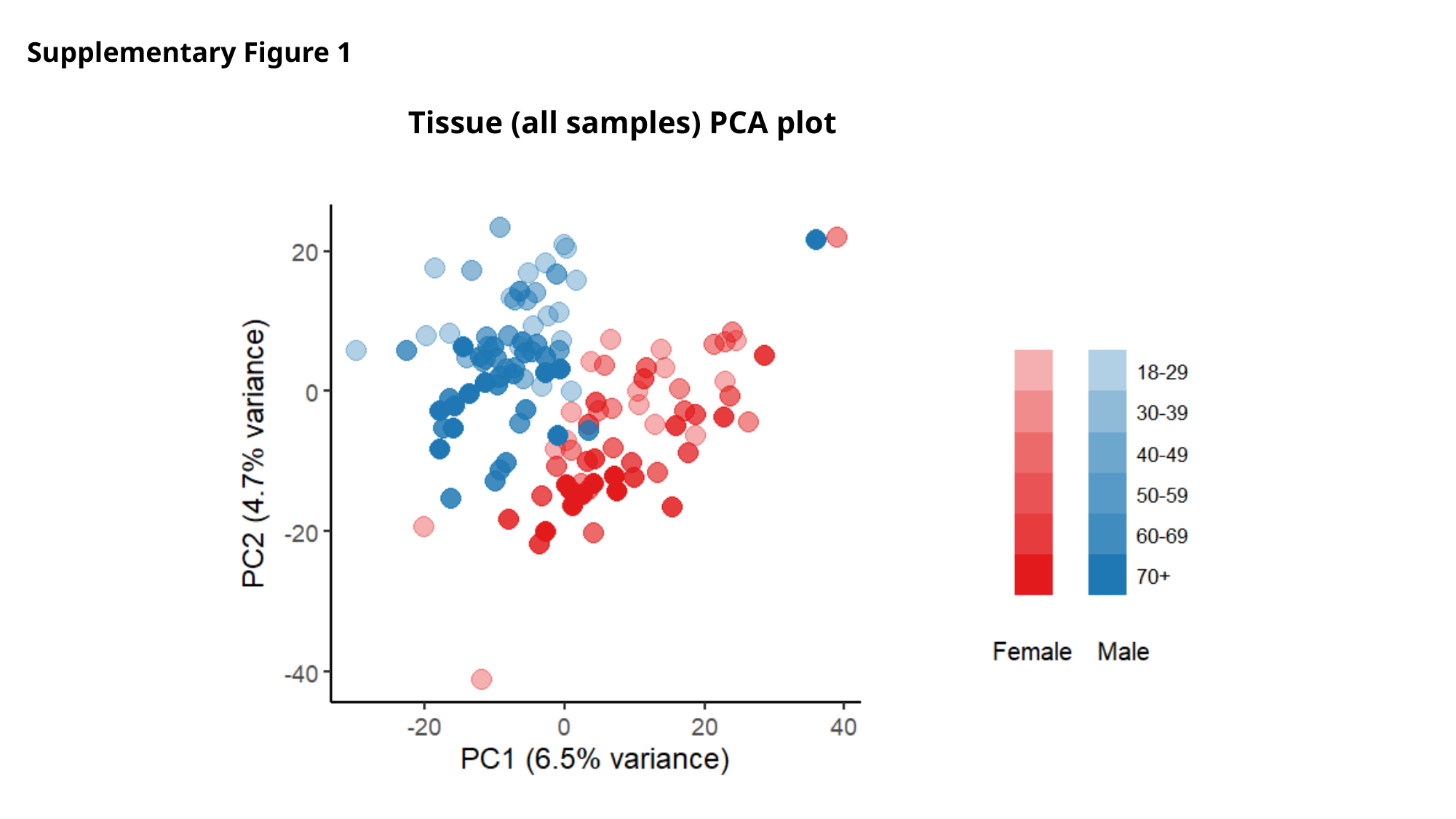

Supplementary Figure 1
Tissue (all samples) PCA plot

### Slide 3
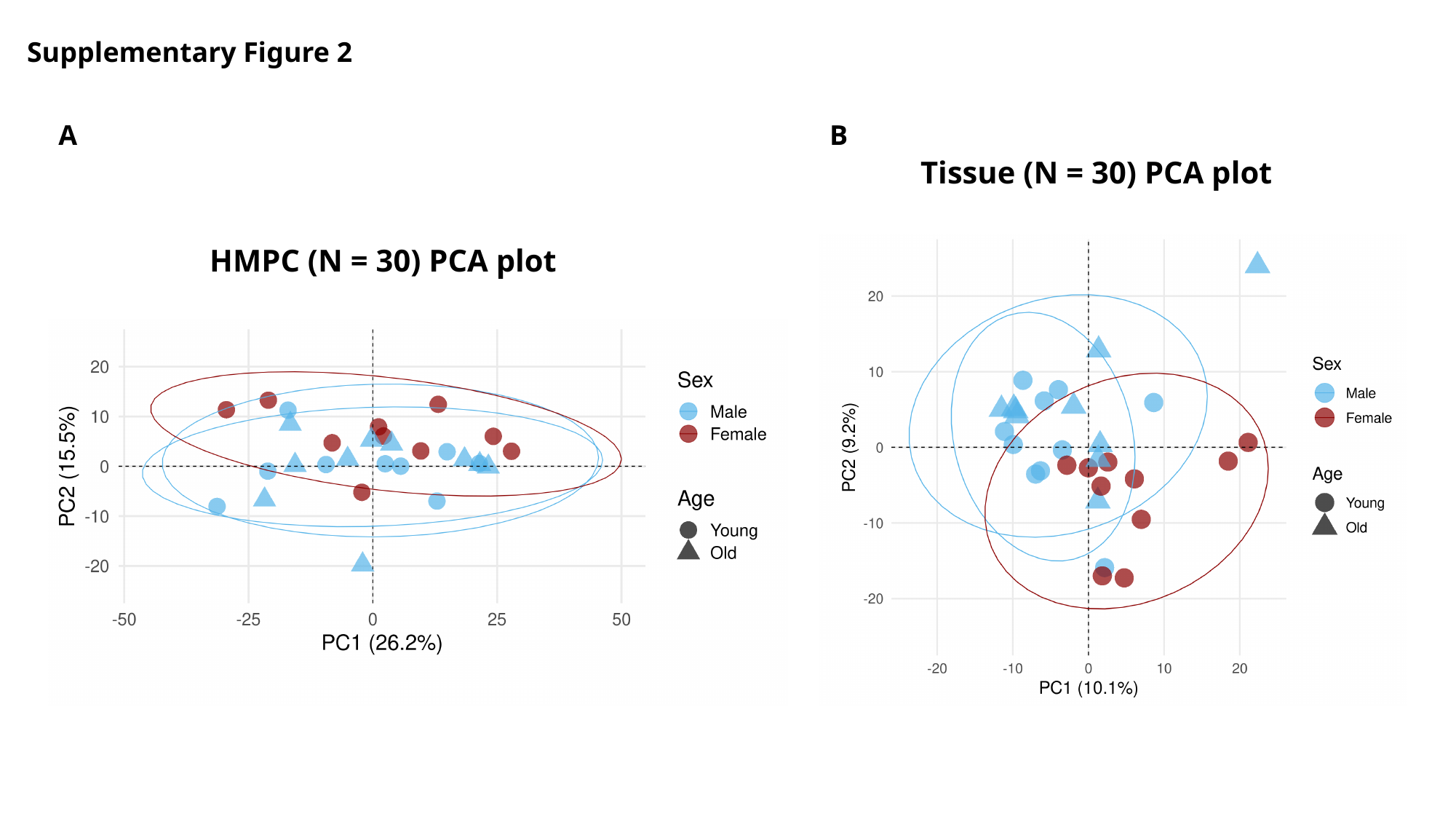

Supplementary Figure 2
A
B
Tissue (N = 30) PCA plot
HMPC (N = 30) PCA plot

### Slide 4
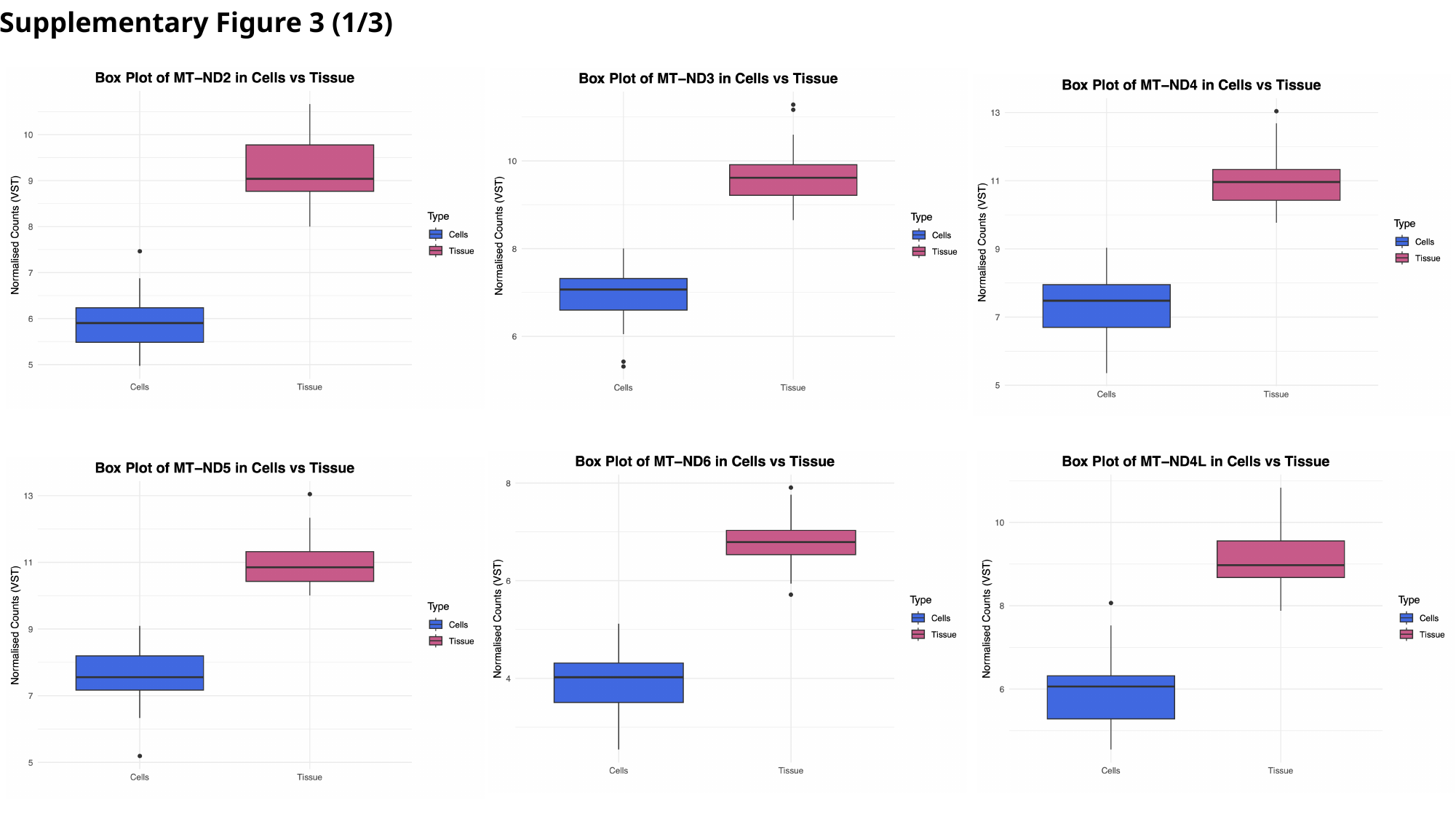

Supplementary Figure 3 (1/3)

### Slide 5
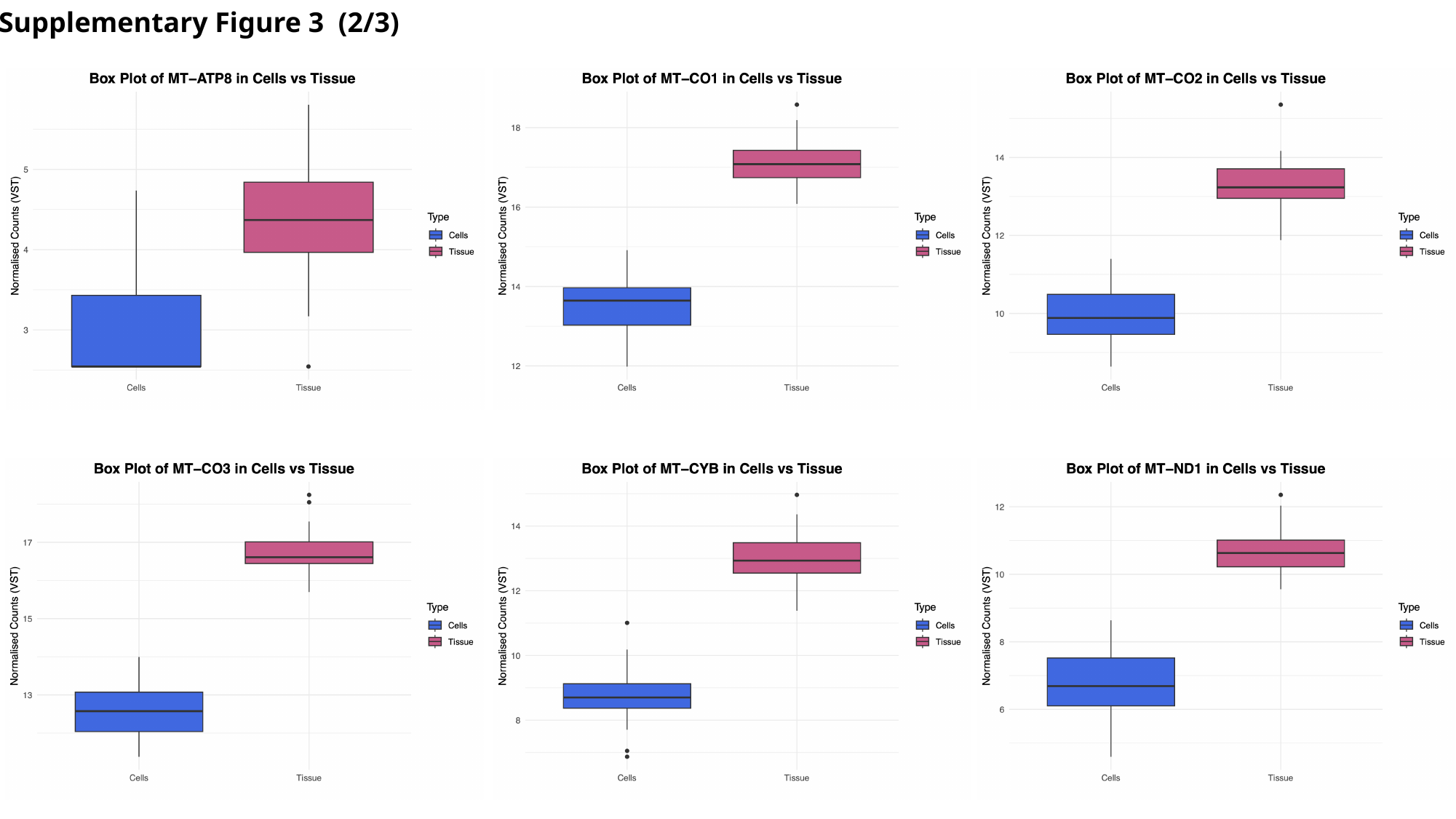

Supplementary Figure 3 (2/3)

### Slide 6
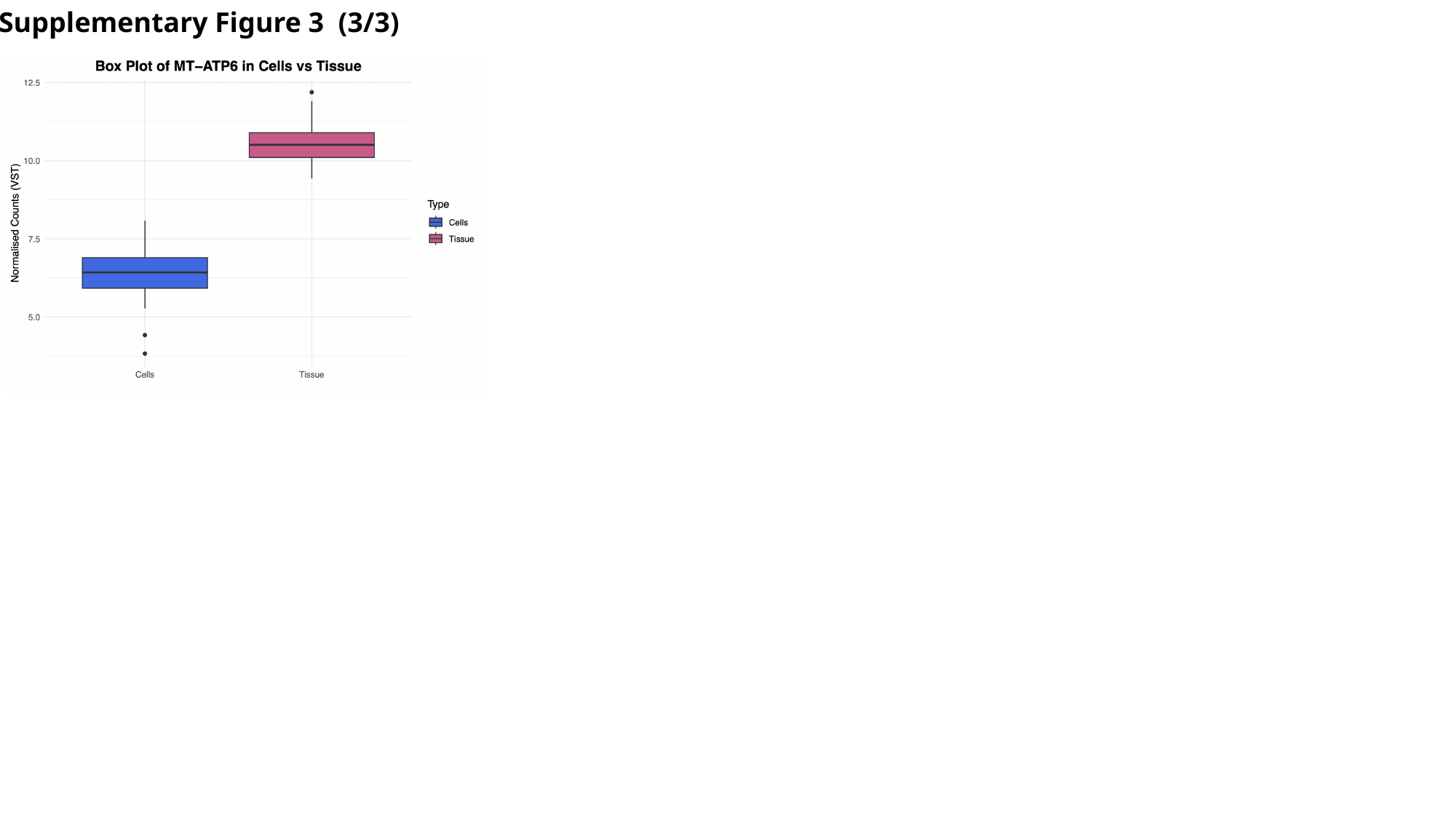

Supplementary Figure 3 (3/3)

### Slide 7
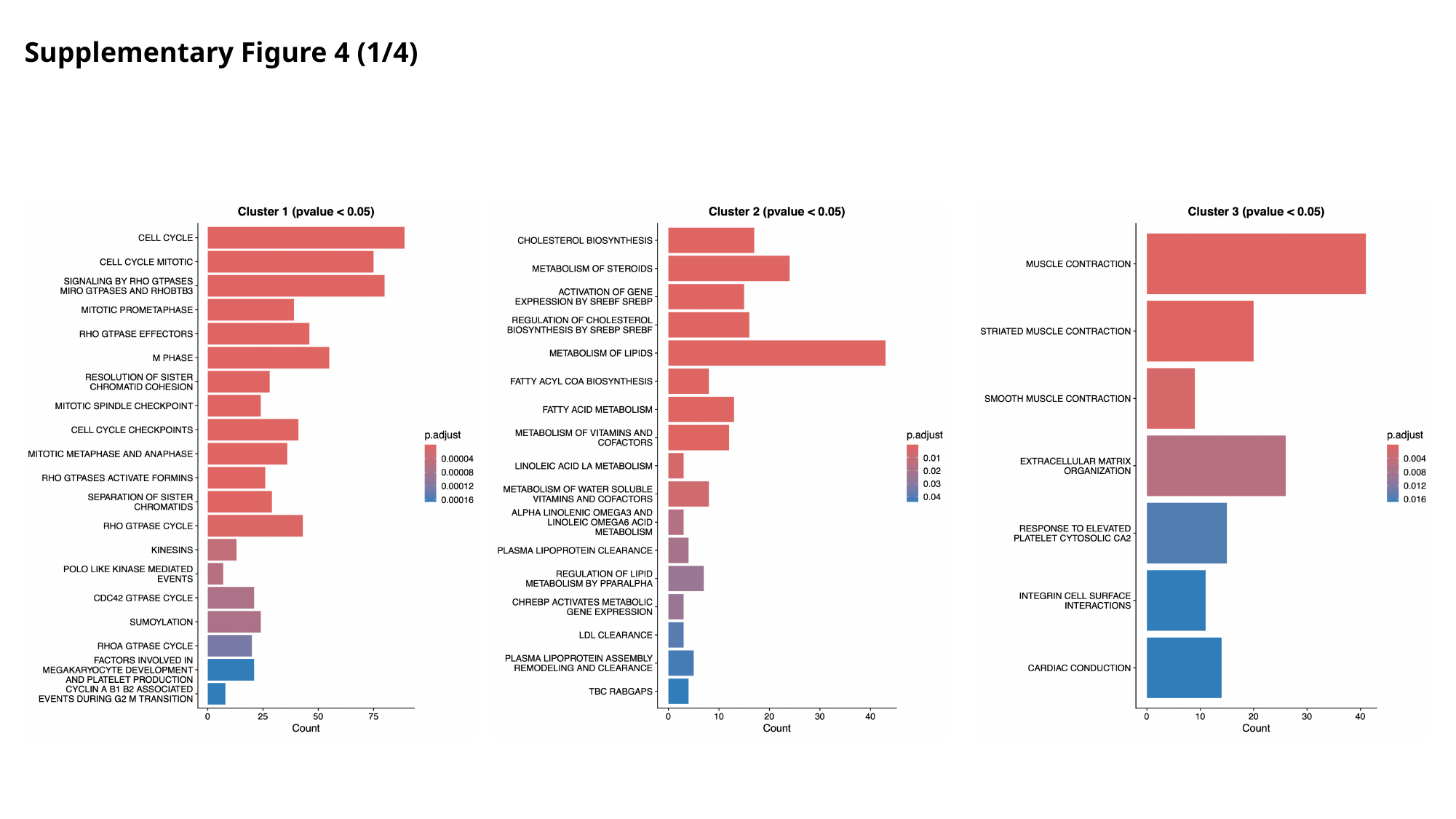

Supplementary Figure 4 (1/4)

### Slide 8
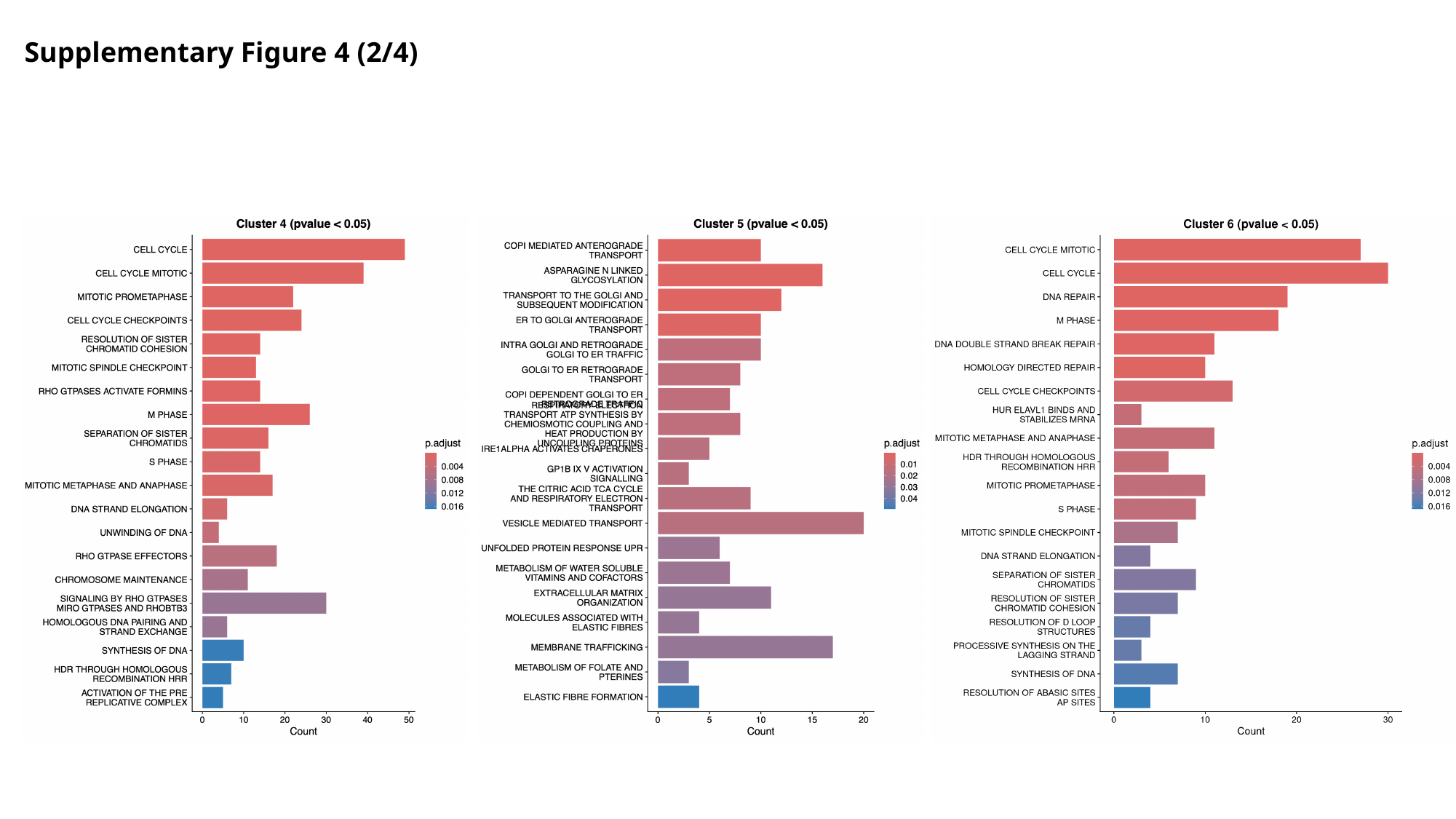

Supplementary Figure 4 (2/4)

### Slide 9
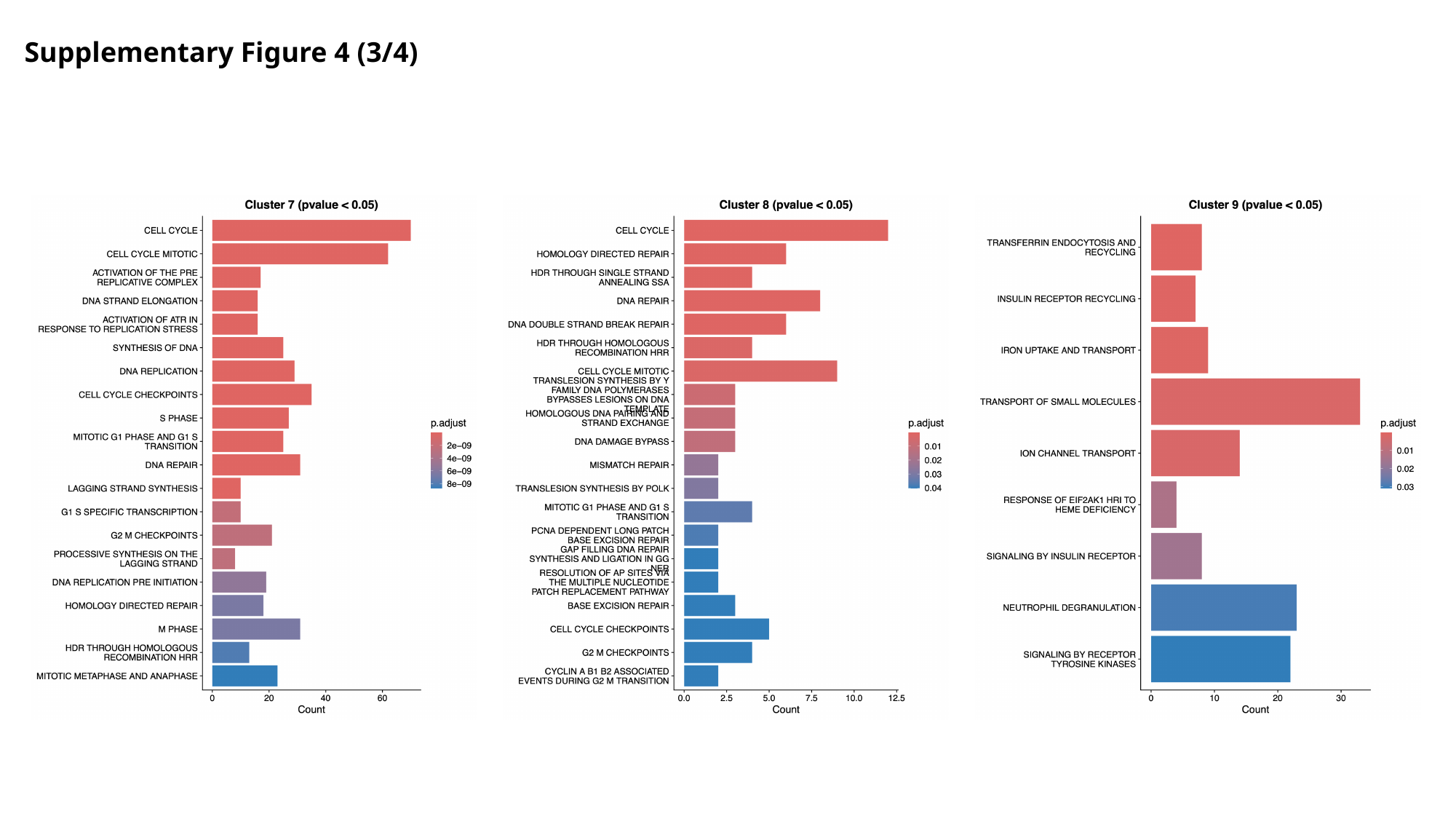

Supplementary Figure 4 (3/4)

### Slide 10
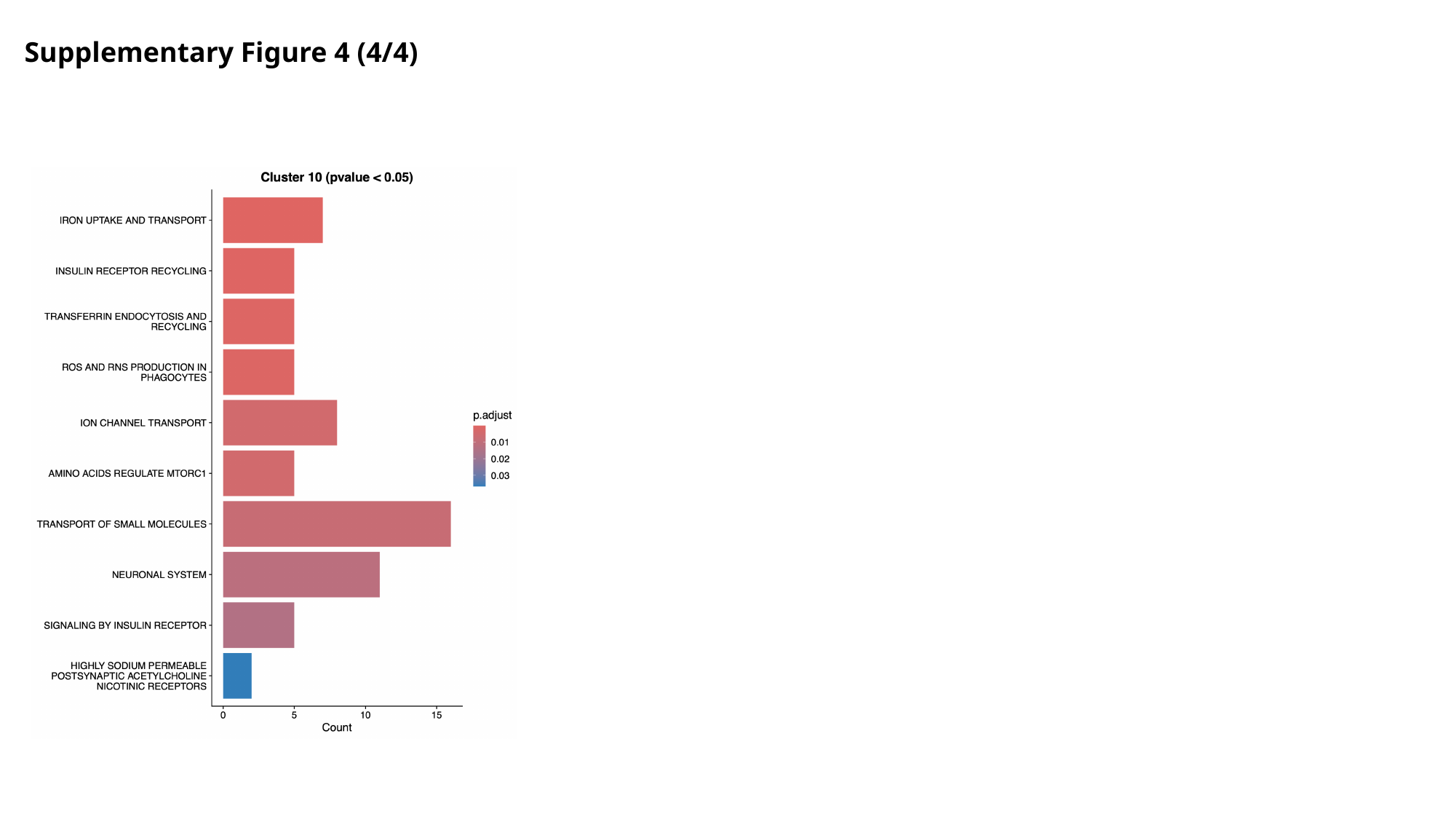

Supplementary Figure 4 (4/4)

### Slide 11
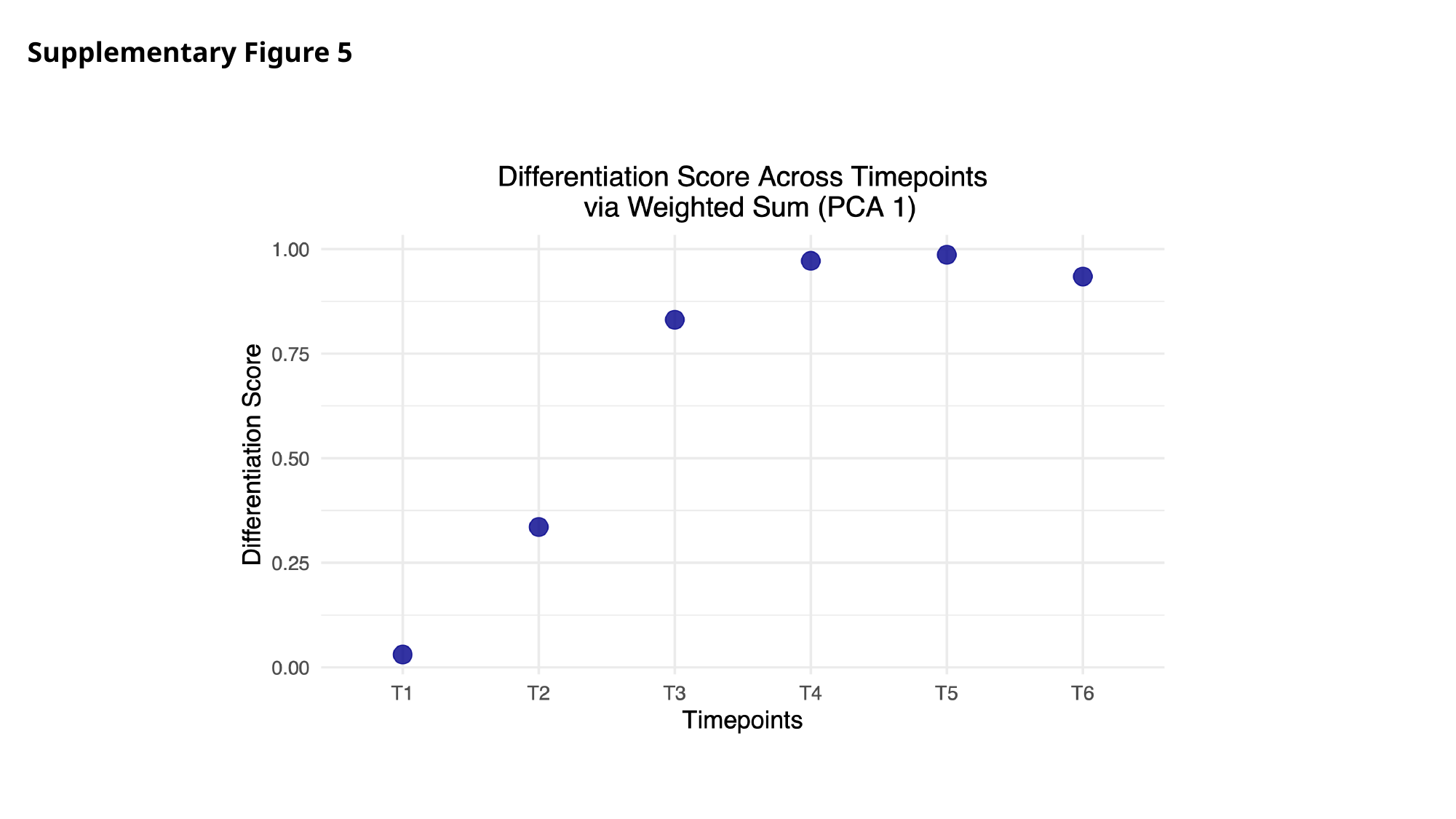

Supplementary Figure 5

### Slide 12
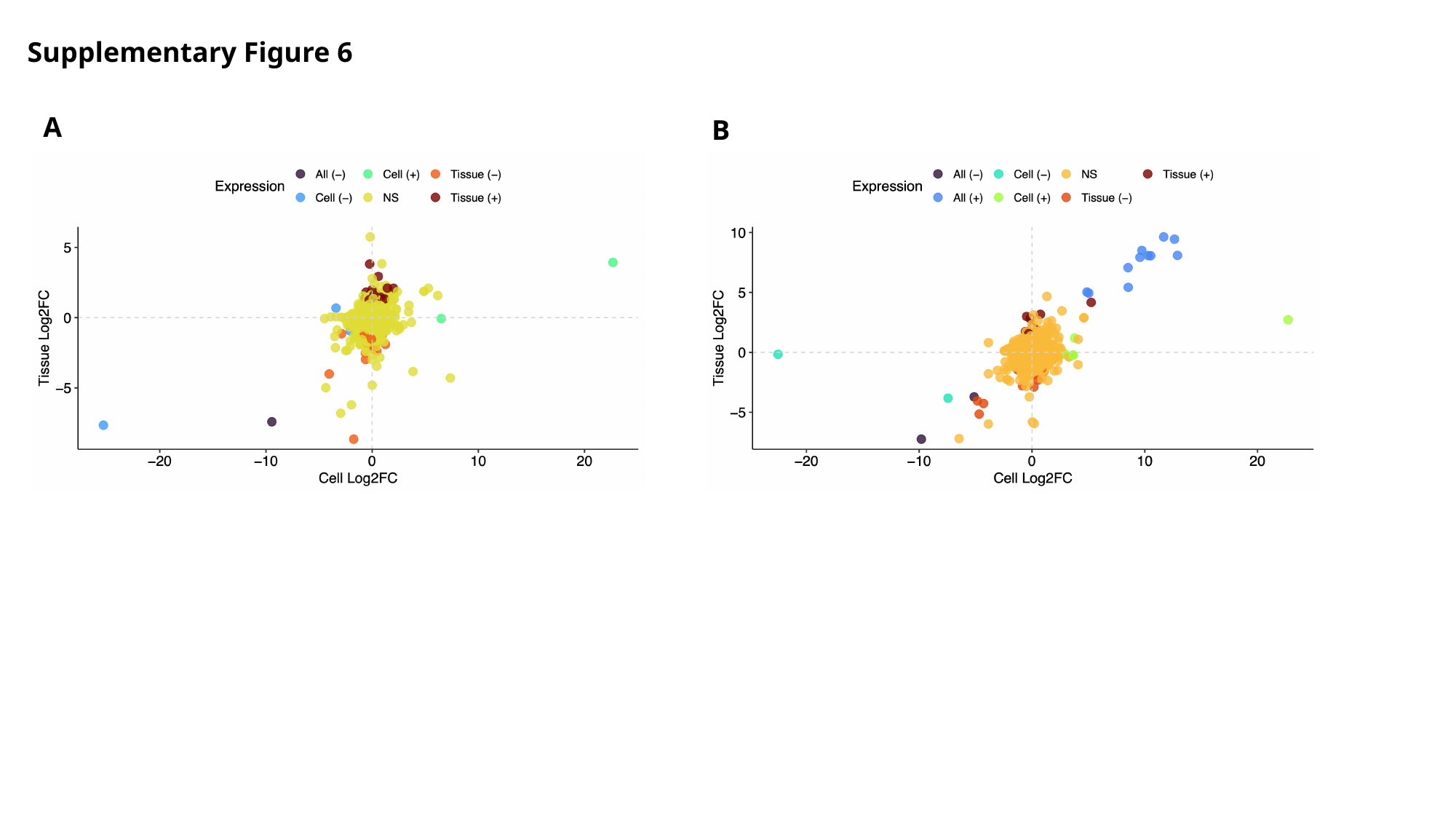

Supplementary Figure 6
A
B
